# TREM2^+^ macrophages accumulate in alveoli of human pulmonary tuberculosis providing a permissive niche for bacterial growth

**DOI:** 10.1101/2025.07.15.664002

**Authors:** Rosane M. B. Teles, Chaouki Benabdessalem, Jonathan Perrie, Cenfu Wei, Julie West, Bruno J. de Andrade Silva, Priscila R. Andrade, Lilah A. Mansky, Prajan Divakar, Linda Fischbacher, Karen Lam, Feiyang Ma, Yiqian Gu, Kimia Rategh, Madeline Brown, Aparna Pillai, Samuel M. French, Emna Romdhane, Mohamed-Ridha Barbouche, Eynav Klechevsky, Marco Colonna, Adrie J. C. Steyn, Steven J. Bensinger, Daniel L. Barber, Soumaya Rammeh, Parambir S. Dulai, Bryan D. Bryson, Matteo Pellegrini, John T. Belisle, Barry R. Bloom, Robert L. Modlin

## Abstract

Pulmonary tuberculosis (TB) exhibits marked spatial heterogeneity, with alveolar pneumonia and organized granulomas frequently coexisting within the same lung. While granulomas have long dominated conceptual models of TB pathogenesis, the immune programs operating within alveolar TB pneumonia in humans remain incompletely defined. Here, we integrate spatial transcriptomics, single-cell RNA sequencing, high-resolution imaging, and functional assays of human lung biopsies to directly compare alveolar pneumonia with adjacent granulomas from the same individuals. We demonstrate that alveolar TB pneumonia is enriched for TREM2⁺ lipid-laden macrophages characterized by lipid metabolic reprogramming, sparse T-cell infiltration, attenuated antimicrobial gene expression, and abundant *Mycobacterium tuberculosis* (*Mtb*) transcripts and antigens. In contrast, neighboring granulomas exhibit organized lymphoid architecture and robust antimicrobial programs. Mechanistically, the mycobacterial virulence lipid phthiocerol dimycocerosate (PDIM) and free mycolic acids induce TREM2 expression and activate TREM2–DAP12 signaling, promoting lipid droplet accumulation, suppressing autophagy, and enhancing intracellular *Mtb* survival in human macrophages. This immunometabolic state is pharmacologically reversible: 1,25-dihydroxyvitamin D₃ downregulates TREM2, restores autophagy, reduces lipid droplets, and limits bacterial viability. Together, these findings define a spatially localized TREM2⁺ foamy macrophage program within alveolar pneumonia that contrasts sharply with adjacent granulomatous immunity, establishing an niche permissive for bacillary persistence and potentially transmission, as well as identifying a tractable host pathway in human TB pathogenesis.

## Introduction

The initial pulmonary infection by *Mycobacterium tuberculosis* (*Mtb*) in human individuals, known as “primary” tuberculosis (TB), involves early pneumonia or granuloma formation^1^. In most cases the infection is contained by host’s immune response, and in some cases can persist for long periods of time. Subsequent reactivation or reinfection, known as “post-primary” TB, initially results in a pneumonia, characterized by the accumulation of foamy macrophages in alveoli^2, 3^. Only after the pneumonia is established, organized granulomas containing predominantly macrophages and lymphocytes develop to further contain the pathogen^2, 4^. The majority of the clinical cases of TB are of the post-primary form.

As early as the 19th century, Laennec recognized that TB could begin as a bronchopneumonia rather than as a granulomatous lesion^5^. Classical pathology studies in the mid-20th century subsequently established that the earliest pulmonary lesion in human TB is an exudative alveolar process^6^. Medlar confirmed these observations in a systematic autopsy series of accidental deaths, identifying early alveolar lesions containing numerous intracellular bacilli but lacking granulomatous organization^7^. Lurie later integrated these human observations with decades of experimental work, demonstrating that bacillary expansion occurs during an initial alveolar phase that precedes granuloma formation^8^. Canetti documented exudative alveolar disease occurring in isolation or adjacent to mature granulomas, with granulomas seldom observed without surrounding alveolar involvement, consistent with an early pneumonia phase that precedes and is later spatially constrained by granulomatous organization^9^. Because early TB pneumonia is rarely diagnosed or biopsied in isolation, lesions containing adjacent alveolar and granulomatous pathology offer a unique opportunity to examine distinct tissue programs within the same host and disease context, without presuming temporal primacy.

The spectacular appearance of granulomas to the naked eye, organized aggregates of macrophages compromising a core and lymphocytes forming a surrounding mantle zone, have long been considered the hallmark lesion of TB^10^. Early microscopic pathology, notably articulated by Virchow, established tuberculous lesions as structured cellular processes rather than amorphous necrosis, laying the foundation for subsequent immunologic interpretations of TB pathology^11^. Accordingly, granulomas have been the predominant focus of immunologic studies of TB.

Human TB granulomas exhibit a highly organized cellular architecture in which CD8^+^ T cells are located in the mantle zone surrounding the core and CD4^+^ T cells are present throughout^12, 13^. The macrophage-rich myeloid core, is composed of CD11b⁺CD11c⁺ and CD68⁺ macrophages, CD163⁺ and CD206⁺ subsets, CD14⁺ monocytes and multinucleated giant cells^14^. B cell rich tertiary lymphoid structures were variably present in or near the mantle zone. Spatial proteomic analyses further demonstrate that granulomas contain concentrated antimicrobial effectors, including cathelicidin and other antimicrobial peptides, as well as reactive oxygen species–associated proteins^15^. Foamy macrophages are a recurrent and defining feature of alveolar TB pathology. Within alveolar pneumonia, these cells accumulate abundant cholesteryl esters and triglycerides^16, 17^ and serve as a nutrient-rich reservoir for *Mtb*^18^. Immunohistology of the alveolar pneumonia in post-primary pulmonary TB has identified that CD163⁺ macrophages are abundant, suggesting macrophage M2-like polarization^1^. In contrast to granulomas, which are enriched for Th1, Th17, and cytotoxic T cells, alveolar lesions dominated by foamy macrophages typically exhibit sparse T-cell infiltration^19, 20, 21^, suggesting a permissive immune environment that may support early bacillary expansion.

Previous studies identified Triggering Receptor Expressed on Myeloid cells 2 (TREM2) as a receptor for non-glycosylated mycobacterial lipids and demonstrated that TREM2 signaling constrains macrophage antimicrobial activation^22^. Subsequent work in human macrophages corroborated and extended these findings, showing that *Mtb* induces TREM2 expression via a STING–type I interferon–dependent pathway and that TREM2 promotes intracellular bacterial survival by suppressing reactive oxygen species and inflammatory cytokine production^23^.

We therefore hypothesized that foamy macrophages in alveolar TB pneumonia represent a TREM2⁺ macrophage state that establishes a permissive alveolar niche for bacillary expansion. Here, using spatial transcriptomics, single-cell RNA sequencing, and high-resolution imaging of human TB lung biopsies obtained during early suspected disease, we define these lesions, map bacterial localization, and test whether mycobacterial lipids and TREM2 signaling drive the foamy macrophage program. Our findings support a model in which TREM2⁺ foamy macrophages constitute a key permissive alveolar niche for *Mtb* survival and potentially transmission.

## Results

We analyzed lung biopsies from four patients with untreated pulmonary TB taken to establish diagnosis, each containing regions of acute pneumonia as well as organized granulomas^9, 21^. Histologically, the pneumonia lesions were evident as clusters of alveoli filled with aggregates of foamy (lipid-rich) macrophages, replacing the normal airspaces (**Figure 1A-D**)^21^. In contrast, nearby granulomas showed the expected architecture: dense central collections of epithelioid macrophages and Langhans multinucleated giant cells, sometimes with central necrosis, surrounded by a mantle of lymphocytes^21^. Langhans-type multinucleated giant cells are easily recognized as large cells characterized by abundant pale eosinophilic cytoplasm and multiple nuclei arranged in a striking peripheral, semicircular or circular pattern along the cell membrane. The pneumonia and granulomas were often adjacent, underscoring that the foamy macrophage pneumonia represents an earlier stage within the same infected region of lung that can give rise to granulomas over time^24^.

**Figure 1.**
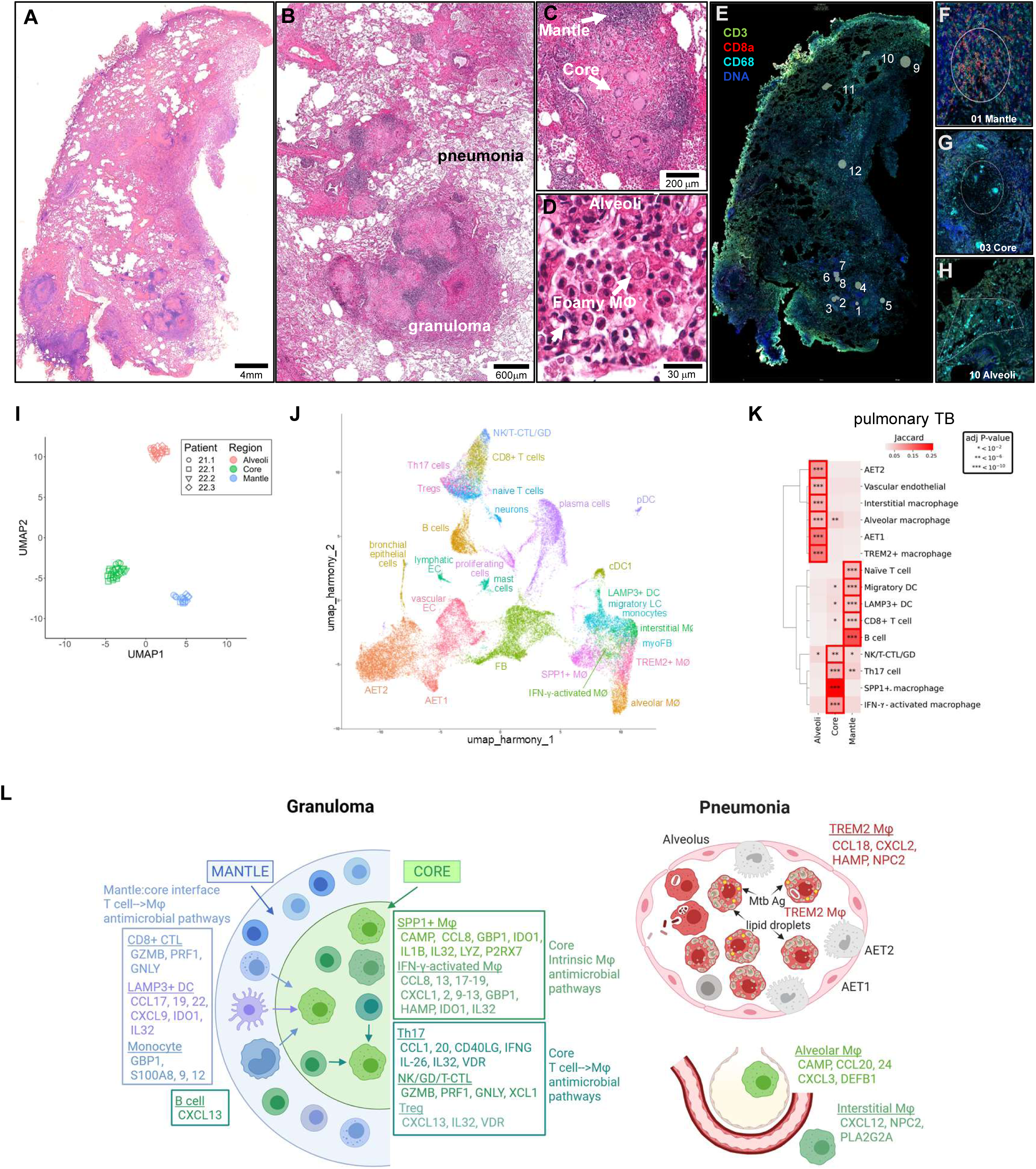
GeoMx spatial profiling of the human transcriptome in pulmonary TB (PTB). (A) Pulmonary TB biopsy specimen from PTB 21.1a, H&E with insets showing (B) higher magnification, (C) Organized granuloma, (D) Pneumonia showing alveoli with foamy macrophages. (E) Regions of interest (ROI), serial section, immunofluorescence, (F) Mantle- ROI 01, (G) Core- ROI 03, (H) Alveoli- ROI 10, (I) UMAP representation of all ROIs. (J) UMAP representation of PTB cell signature identified by scRNA-seq. (K) Jaccard overlap of the GeoMx spatial profiling data with the PTB scRNA-seq data. (L) Distribution of antimicrobial genes in PTB identified by integrating the GeoMx spatial profiling data with the scRNA-seq

In this study, we performed detail analysis of all four specimens, performing GeoMx spatial transcriptomic profiling, scRNA-seq, CosMx spatial proteomics and immunohistology. Using GeoMx spatial transcriptomic profiling, we quantitatively compared gene expression in micro-dissected regions of interest from alveolar pneumonia, granuloma cores, and granuloma mantles (**Figure 1E–H, Supplemental Table 1, Supplemental Figure 1**). The images for each ROI and their location in mantle, core, alveoli are shown in **Supplemental Figure 2.** Unsupervised clustering of the transcriptomes clearly segregated regions of interest by their spatial location (alveoli vs. core vs. mantle), indicating distinct molecular signatures for each compartment (**Figure 1I**). Differential expression analysis revealed that the alveolar pneumonia regions were enriched for genes involved in lipid metabolism and surfactant handling, consistent with the presence of foamy macrophages within the alveolar pneumonia and Type II pneumocytes. Notably, the alveoli showed high expression of mRNAs encoding apolipoproteins and lipid-processing enzymes (e.g. *APOE*, *APOC1*, *LIPA*), as well as scavenger receptors like *CD36* and *CD163* (**Supplemental Figure 3A, B**), which are known markers of alternatively activated, lipid-associated macrophages^14, 25^. Alveolar regions also contained transcripts from alveolar type II epithelial cells (*SFTPA2, SFTPC* encoding surfactant proteins)^25^, reflecting epithelial cells lining alveoli. In stark contrast, the granuloma core regions over-expressed a suite of pro-inflammatory and antimicrobial genes, including *IL1B, CYP27B1, MT2A* and *CYBB*^14^ (**Supplemental Figure 3A, C**). These core signatures are indicative of classically activated (“M1-like”) macrophages and Th1/Th17-type T cells working to control infection. The granuloma mantles were enriched for B and T lymphocyte genes (e.g. *IGHG, CD3E, CD8A*), consistent with lymphocyte clustering around granulomas^13^ (**Supplemental Figure 3B, C**). By computing the set intersection of genes differentially expressed in one region versus the other two, we identified signature genes for each region: alveoli=559, core=414, and mantle=803 (**Supplemental Table 3**).

### What are the cell types and their gene signatures that comprise human pulmonary TB?

We used scRNA-seq on the lung tissue to identify specific cell populations present in these lesions. Clustering of ∼45,000 cells in total across the four patients (each patient contributed between 8,000 and 15,000 cells; data were integrated for joint analysis) yielded a total of 27 cell types, which were identified by comparing to the scRNA-seq cluster signatures to known cell signatures including major cell types^26, 27^, a human lung atlas^28^, leprosy lesions^29^ and using Enrichr (**Figure 1J, Supplementary Table 4, Supplemental Figure 4A**). Notably, among five distinct macrophage clusters, we identified a cluster corresponding to “TREM2⁺ lipid-associated macrophages” as well as “IFN-γ-activated macrophages”, “SPP1^+^ macrophages”, “interstitial macrophages”, “alveolar macrophages”. The T cell populations included: CD8^+^ T cells, naïve (or resting) T cells, Th17 cells (with features of Th1 cells and MAITs), NK/T-CTL/ γδ T cells (reflecting cells with expression of genes in the cytotoxic pathways including NK cells, tri-cytolytic T cells (T-CTL) and γδ T cells), and Tregs. These populations were stable across the four different patients (**Supplementary Figure 4B**).

### What are the differences in frequency of cell types between TB pneumonia and organized granulomas?

By integrating scRNA-seq cluster–defined gene signatures with spatial transcriptomic data, we assigned predominant cell types to each tissue region using Jaccard similarity indices, which quantify the overlap between cluster marker gene sets and region-enriched genes. The alveolar pneumonia regions showed the strongest enrichment for the TREM2⁺ macrophage signature (adjusted p < 10⁻¹⁰) (**Figure 1K**), indicating that the lipid-laden macrophages in the alveoli correspond to the TREM2-expressing, lipid-associated macrophages. The alveolar regions also overlapped with signatures of “alveolar macrophages” and “interstitial macrophages”, whereas the granuloma core overlapped most with “SPP1⁺ macrophages” and IFN-γ-activated macrophages. The location of “TREM2⁺ macrophages” in the alveoli is consistent with the location of foamy macrophages in alveoli. The granuloma core contained “SPP1^+^ macrophages”, “IFN-γ-activated macrophages” and “Th17 cells”. The granuloma mantle zone contained “B cells”, “plasma cells”, “CD8^+^ T cells” and “LAMP3^+^ DC”.

Overlap of the GeoMx spatial signature genes from each region of the PTB samples with cell type markers from a scRNA-seq healthy lung atlas^28^ confirmed the cell locations inferred from the pulmonary TB scRNA-seq data, also revealing that the alveoli had strong overlap with gene signatures from “AET1” and “AET2”, consistent with the presence of alveolar epithelial cells (**Supplemental Figure 5A**). The GeoMx PTB spatial signatures were also compared to our previous leprosy lesion scRNA-seq dataset^29^, as leprosy represents a mycobacterial disease with a spectrum of immune responses that correlate with clinical presentation. This comparison demonstrated that macrophages populating the alveolar pneumonia regions share a transcriptional program with “TREM2⁺ macrophages” in the progressive lepromatous form of the disease (**Supplemental Figure 5B**). This transcriptional similarity is independent of location and reinforces that the foamy macrophages in TB alveoli correspond to a conserved TREM2-driven, lipid-associated macrophage state.

We further assessed the different host responses in the granuloma and pneumonia by functional pathway analysis of the pulmonary TB DGE signatures from the GeoMx spatial profiling data. The alveoli DGEs were enriched for lipid metabolism, localization and transport, typical for TREM2⁺ macrophages, as well as surfactant metabolism (**Supplemental Figure 6A**). The core DGEs were enriched for innate immune responses including responses to bacterium while the mantle DGEs were enriched for T and B cell pathways, consistent with the histology of this region and the overlap with leprosy scRNA-seq marker sets. Similar analysis of the pulmonary TB scRNA-seq myeloid cell gene signatures indicated that IFN-γ-activated macrophages and SPP1^+^ macrophages were enriched for “defense response to bacterium” and “antimicrobial humoral immune response mediated by antimicrobial peptide” (**Supplemental Figure 6B**). The TREM2⁺ macrophage and alveolar macrophages were enriched in lipid metabolism pathways.

### Do TREM2⁺ macrophages preferentially localize to the alveoli in TB pneumonia?

To further clarify the location of TREM2⁺ macrophages in pulmonary TB, we derived a gene signature using a consensus set of 34 TREM2⁺macrophage related scRNA-seq gene signatures from 21 publications (**Supplemental Table 7**), ranking the frequency of genes in these signatures (**Supplemental Tables 8 and 9**). Overlap with the pulmonary TB DGEs revealed that the twelve most highly ranked TREM2 macrophage signature genes had a higher overlap with alveoli than core, with the ratio of the Jaccard indices greater than 5, for genes found in ≥22 signatures (**Supplementary Figure 8A, B**). The expression of eight genes, *LGMN*, *CTSD*, *TREM2*, *NPC2*, *APOE*, *C1QB*, *GRN* and *C1QC* was greater in the alveoli vs. core, whereas *C1QA* expression was almost equal and *CTSB* was greater in the core, with *MS4A6E* and *FTL* not detected in the GeoMx data (**Supplemental Figure 8C**). We examined the TREM2⁺ macrophage scRNA-seq signature genes, which significantly overlapped with the gene expression in the alveoli in the GeoMx data (q<10^−10^). Eleven of the twelve TREM2⁺ macrophage genes listed above were indeed present in the TREM2⁺ macrophage scRNA-seq cluster from our scRNA-seq data (all except MS4A6E).

Given that TREM2⁺ macrophages are characterized by expression of lipid metabolism genes^30^, we mined the TREM2⁺ macrophage scRNA-seq data, finding enrichment for genes for lipid metabolism pathway genes characteristic of lipid localization, lipid metabolism, lipid signaling, response to lipids, cholesterol transport and cholesterol efflux (**Supplemental Figure 8D, Supplemental Table 10**). Of these, *TREM2*, *APOE*, *CTSD*, *NPC2*, *PLIN2*, *LIPA*, *PSAP*, *CD81*, *APOC1* and *HEXB* were present in 17 of the 34 TREM2⁺ macrophage gene signatures. Together, these data demonstrate that the TREM2⁺ macrophages expressing lipid metabolism genes are predominantly located in the alveoli of the pulmonary TB pneumonia.

### Do antimicrobial programs differ between pneumonia and granulomas?

Spatial and single-cell analyses revealed striking compartmentalization of antimicrobial responses within the same lungs. The TB pneumonia in alveoli was dominated by lipid-associated macrophages and showed minimal expression of canonical TB antimicrobial products, whereas organized granulomas exhibited a concentrated antimicrobial inflammatory profile. Using a curated panel of 52 antimicrobial genes from the Antimicrobial Peptide Database 3^31^ and key mycobacterial host-defense genes^29, 32^ (**Supplemental Table 5**), we detected markedly fewer antimicrobial transcripts in alveolar regions of interest compared with granuloma cores (3/52 vs. 15/52, q < 10⁻⁶), despite the bacterial burden in pneumonia. By region, eight antimicrobial genes were detected in granuloma cores (adj. p = 6.78×10⁻⁵), five in the mantle (ns), and three in alveoli (ns) (**Supplemental Figure 7A-B; Supplemental Table 6**). Direct comparison showed a significant enrichment of antimicrobial transcripts in granuloma cores relative to alveoli (15 genes, adj. p = 3.72×10⁻⁷).

Of the 52 antimicrobial genes, 40 were expressed across T cell and myeloid subsets in scRNA-seq of human pulmonary TB (**Supplemental Figure 7C-D; Figure 1L**). The granuloma cores showed the broadest antimicrobial profile: IFN-γ-activated macrophages expressed fifteen antimicrobial genes, SPP1⁺ macrophages eight, Th17 cells seven, and NK/T-CTL/γδ T cells four. In the mantle, CD8⁺ T cells expressed three antimicrobial genes, LAMP3⁺ dendritic cells six, and monocyte-like cells four. Notably, several of these encode proteins with direct antimycobacterial activity: *CAMP* (cathelicidin) in SPP1⁺ and alveolar macrophages^33^; *GNLY* (granulysin) in CD8⁺ T and NK/T-CTL/γδ T cells^29, 34, 35^; *IL26* in Th17 cells^36^; and *S100A12* in monocytes^37^. This dichotomy highlights the functional specialization of the lesions: granulomas are immunologically organized for containment, whereas the pneumonia would appear to be relatively permissive to *Mtb*.

### What are the differences in the frequency of cell types as defined by specific protein determinants between TB pneumonia and the organized granulomas?

We validated these findings at the protein level using high-resolution spatial proteomics (NanoString CosMx) and immunohistochemistry on serial tissue sections. Staining for macrophage and T cell markers recapitulated the lesion architecture: CD68⁺ macrophages and CD68⁺Langhans multinucleated giant cells concentrated in granuloma cores, CD3⁺ T cells and CD20⁺ B cells forming a mantle around granulomas (**Figure 2A, B**). The Langhans multinucleated giant cells are strikingly evident in the granuloma core as large ovoid or round cells, 20-50 µM in diameter by their intense blue CD68⁺staining. The ratios and aggregation of T cells and B cells in the mantle zone varied among granulomas (**Supplemental Figure 9**). Strikingly, CD163⁺ macrophages filled the alveolar spaces of the pneumonia (**Figure 2A, C**). CD163 is a scavenger receptor induced in M2-polarized and foamy macrophages, and its presence correlated with the foamy macrophages in alveoli. Within alveoli, macrophages varied in CD68/CD163 expression, either CD163⁺-predominant (**Figure 2D**) or CD68⁺CD163⁺ co-expressing (**Figure 2E**), yet were uniformly foamy on H&E. In contrast, CD3⁺ T cells were rarely present (**Figure 2F and Supplemental Figure 10**).

**Figure 2.**
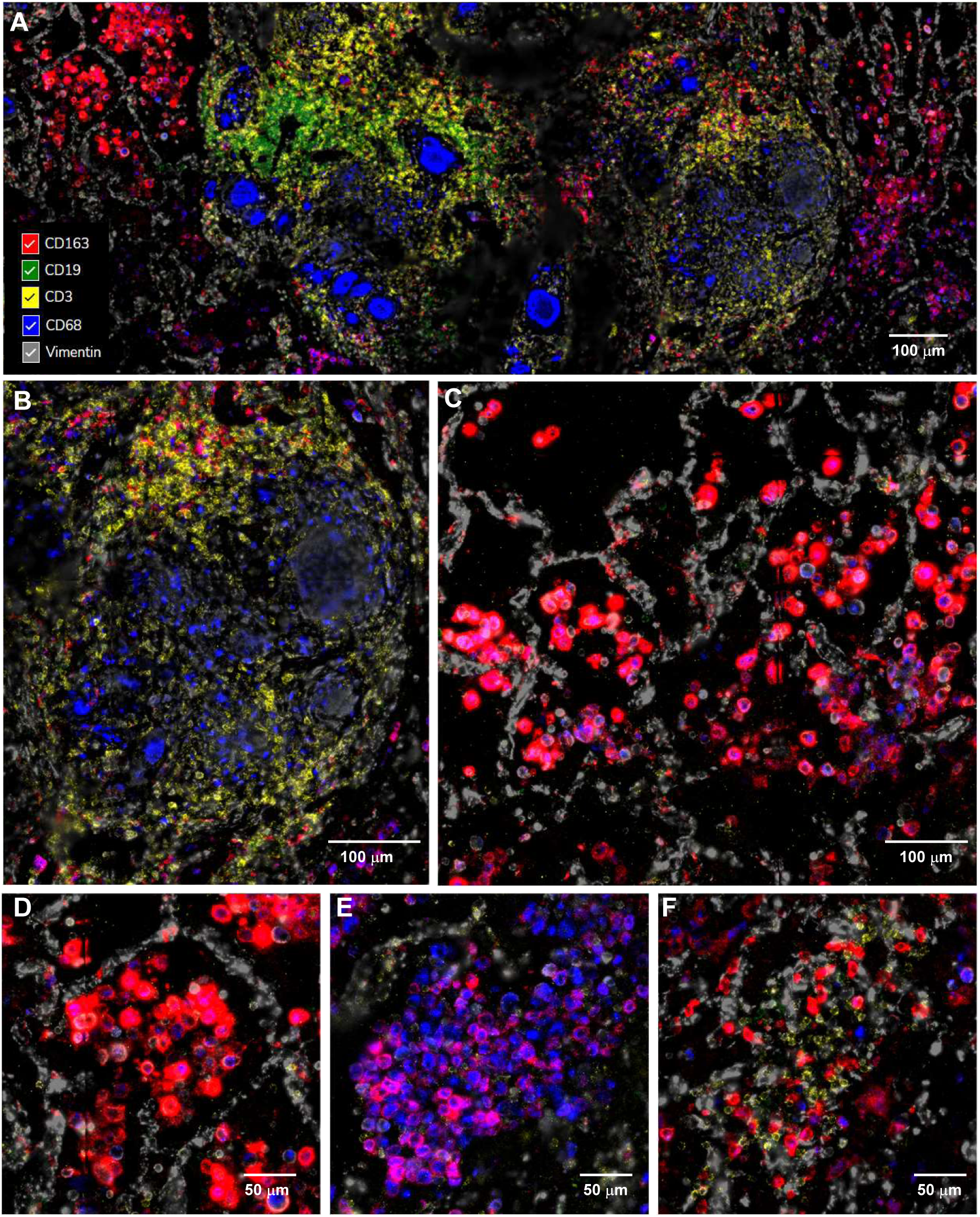
Immunofluorescence images obtained using the CosMx spatial profiling with the human protein panel, illustrating the spatial expression of immune cell markers in active pulmonary tuberculosis (TB). (A) Overview of immune cell distribution within the active pulmonary TB lesion. The granuloma core contains CD68^+^ macrophages (blue) and CD3 ^+^ T cells (yellow). The granuloma mantle contains CD3 ^+^ T cells (yellow) and CD19 ^+^ B cells (green). The alveoli contain prominent CD163 ^+^ macrophages (red), some with CD68 expression (blue). Vimentin (gray) is present on many cells in the lesions but particularly highlights the alveolar structure by staining alveolar epithelial cells. (B) Localization of immune cells within granulomas. (C–F) Distribution of immune cells in alveolar regions showing heterogeneity of macrophage phenotype and variability in the presence or absence of T cells. Scale bars: 100 µm (low magnification) and 50 µm (high magnification).

The CD163⁺ foamy macrophages in alveoli were often found in close proximity to granulomas, for instance, surrounding small nascent granulomas or in alveoli adjacent to a granuloma mantle (**Supplemental Figure 9A, 11**). This spatial relationship supports the concept that some, if not most granulomas arise physically within, the initial foamy macrophage pneumonia. Another hallmark feature of pulmonary TB, bronchiole obstruction, a mechanism associated with spread of the infection^4^, was visualized with surrounding lymphocytic infiltrate containing T and B cells, necrosis, and obliteration of the bronchiolar epithelium such that the airway was occluded by necrotic granulomas (**Supplemental Figure 12**). Three-dimensional microCT studies of human TB lungs reveal branched, airway-connected necrotic granulomas and bronchi filled with caseous material, indicating that pneumonia and granulomatous lesions commonly coexist and may arise through bronchogenic spread rather than a simple linear progression^38^.

By standard immunohistology, we detected CD3^+^ and CD4^+^ lymphocytes throughout the granuloma, with CD8^+^ and CD20^+^ lymphocytes predominantly localized to the mantle zone (**Supplemental Figure 13A-D**). CD68^+^ macrophages and Langhans giant cells were confirmed to be in the core of the granuloma, with CD163 identifying macrophages in alveoli (**Supplemental Figure 13E, F, and Supplemental Figure 14**).

### What are the differences in localization and phenotype of TREM2⁺ macrophages in the TB alveolar pneumonia compared to granulomas?

Immunostaining for TREM2 itself revealed a striking pattern: TREM2 protein was highly expressed on the foamy macrophages in the alveolar pneumonia, with strong membrane staining on those cells, whereas granuloma macrophages and giant cells had only weaker or patchy TREM2 (**Figure 3A, Supplemental Figure 15 A-D**). These results are consistent with Shang et al who focused on TREM2 staining in the multinucleated giants cells^39^. The macrophages in the alveoli expressed the lipid droplet marker perilipin-2 (PLIN2), CD68, APOE, LIPA and SPP1 (**Figure 3A-B**), indicating they contained abundant intracellular lipid droplets as shown by perilipin-2 (PLIN2) staining (a lipid droplet coat protein)^40, 41, 42^. PLIN2 immunofluorescence exhibited a punctate pattern throughout the macrophages in the alveolar pneumonia, confirming their foamy, lipid-loaded nature (**Supplemental Figure 16**). In sum, these data identify a population of TREM2⁺, CD163⁺, PLIN2⁺ foamy macrophages specifically localized to TB alveolar pneumonia, distinct from the macrophages inside granulomas.

**Figure 3:**
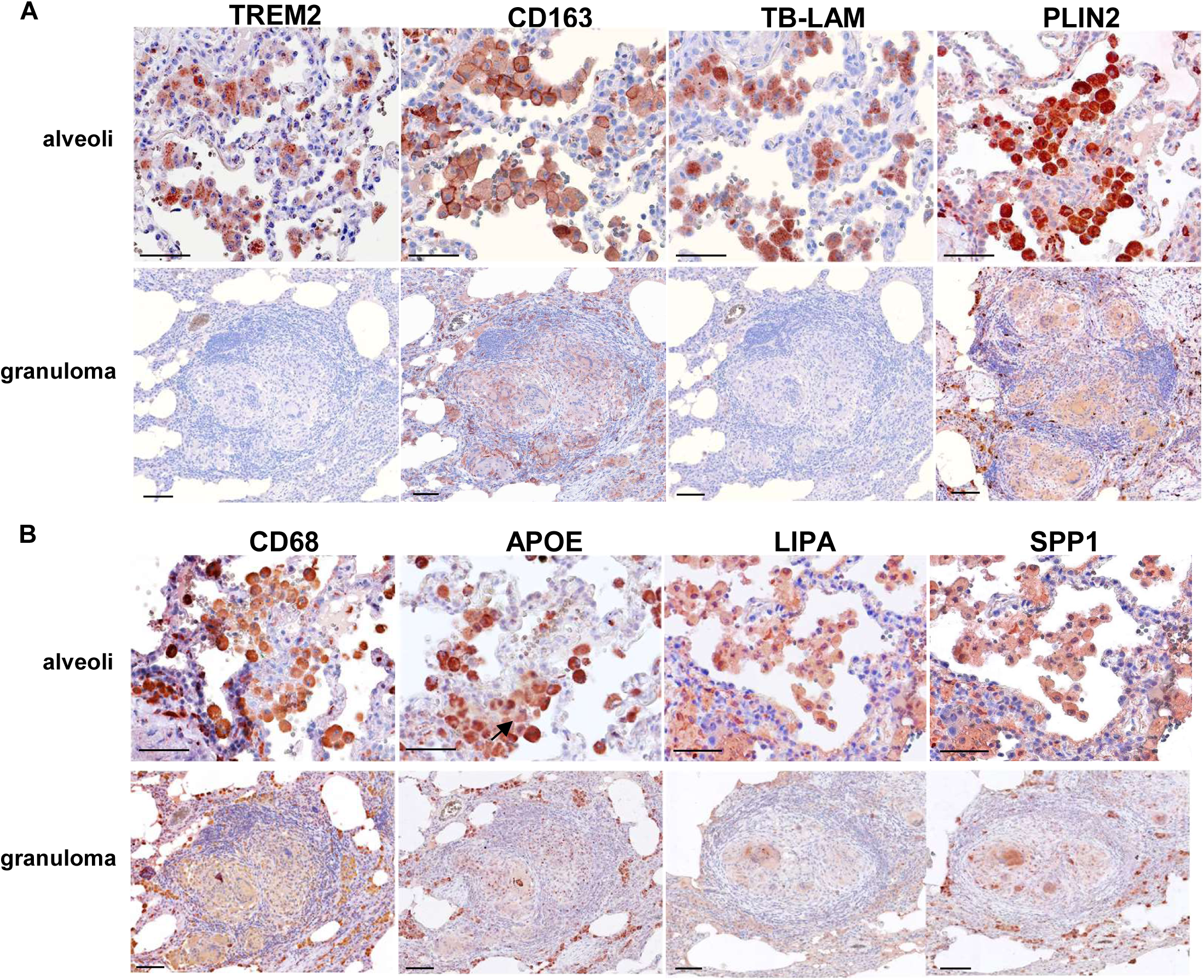
Immunohistochemical analysis of pulmonary tuberculosis (PTB) lung tissue demonstrates enrichment of TREM2-associated macrophage markers within areas of alveolar pneumonia compared with granulomatous regions. (A) TREM2, CD163, and PLIN2, together with *Mtb* lipoarabinomannan (TB-LAM), are predominantly localized in alveolar pneumonia lesions relative to granulomas. (B) CD68, APOE, LIPA, and SPP1 show strong staining in alveolar regions compared with the granuloma core. Images are representative of four independent PTB samples. Scale bars: 40 μm.

We confirmed that the macrophages in the alveolar pneumonia co-expressed CD163 with TREM2 (**Figure 4A**). By immunofluorescence, TREM2 co-localized with LAM in macrophages within the alveolar pneumonia (**Figure 4B**)^43^, indicating that the very TREM2⁺ foamy macrophages are the cells harboring mycobacterial lipid antigens. Moreover, PLIN2⁺ lipid droplets were often adjacent to or overlapping with LAM signals (**4C**), consistent with the known interaction of bacilli and bacterial lipids with host lipid stores^16, 18^.

**Figure 4.**
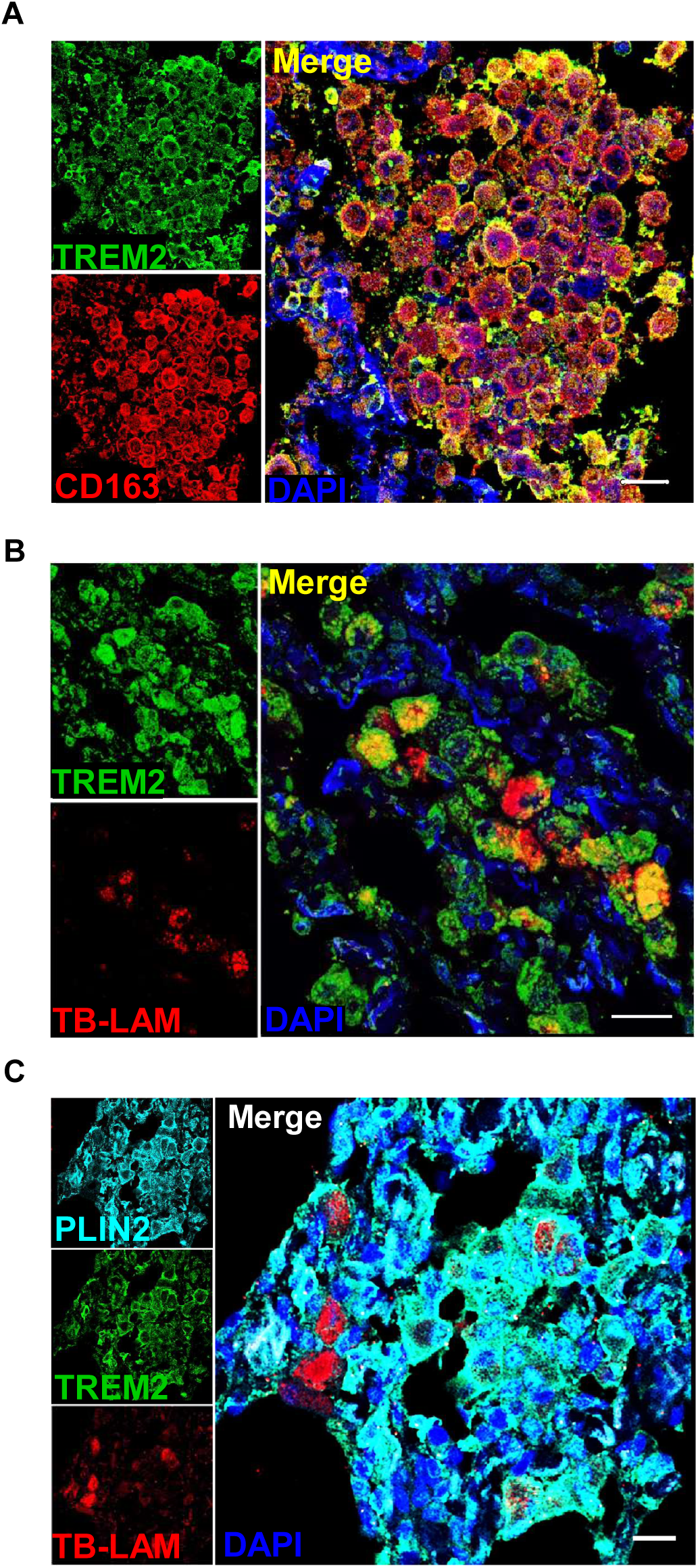
TREM2⁺ macrophages associate with TB-LAM and lipid droplets in PTB lesions. Representative immunofluorescence images showing (A) co-expression of TREM2 (green) and CD163 (red), (B) colocalization of TB-LAM (red) with TREM2⁺ macrophages (green), and (C) co-expression of TREM2 (green), PLIN2 (cyan), and TB-LAM (red); nuclei are counterstained with DAPI (blue). Images are representative of four independent pulmonary tuberculosis (PTB) specimens. Scale bars, 20 µm.

Staining for the major cell wall lipoglycan LAM and for purified protein derivative (PPD, a mix of mycobacterial proteins) revealed the strongest antigen signals in the macrophages within the alveolar pneumonia, whereas the cores of the granulomas showed no signal for LAM and weaker staining for PPD (**Figure 5A, B, C**). Antigen deposits appeared granular and co-localized with foamy macrophages. Robert Hunter previously documented PPD accumulation in the foamy alveolar macrophages^44^. PPD has also been strongly detected in necrotic regions of pulmonary TB, although regions of pneumonia were not clearly investigated^45^.

**Figure 5:**
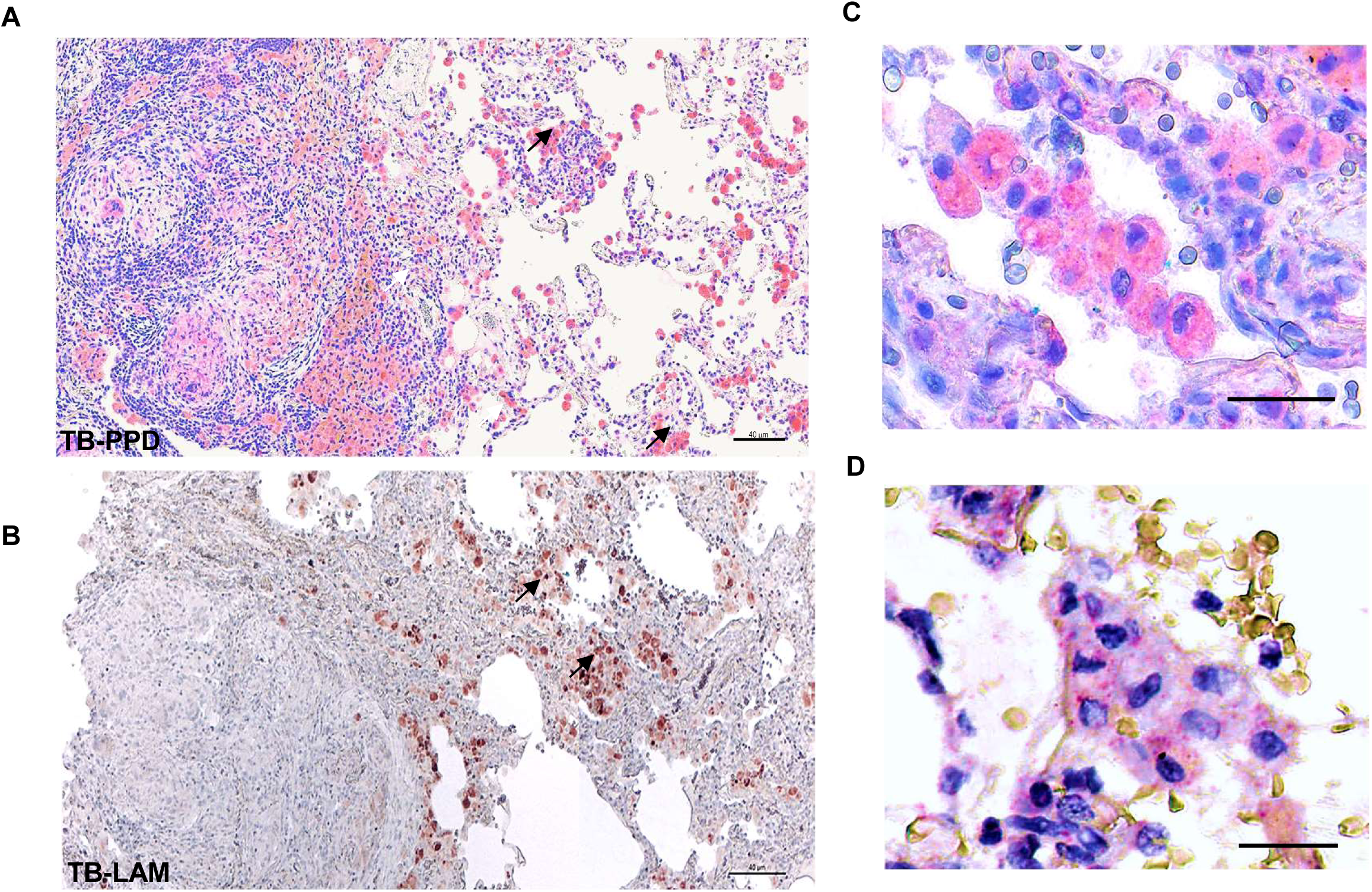
In situ identification of *M. tuberculosis (Mtb)* in PTB tissue shows that (A) TB-PPD (red-pink) is expressed in both alveoli (black arrows) and the granuloma core (white arrows). (B) TB-LAM (red-brown) is expressed in alveoli (black arrows). (C) TB-PPD is strongly expressed in alveolar foamy cells (E) *Mtb* RNA is observed as red puncta present in foamy macrophages within alveoli by in situ hybridization (RNAscope), counterstained with hematoxylin. The brown-staining cells represent erythrocytes. Scale bars: 40 μm (A and B) and 20 μm (C and D).

### Do TREM2⁺ foamy macrophages in TB pneumonia harbor *Mtb* mRNAs, indicating bacterial persistence?

A critical question was whether foamy macrophages within the alveolar pneumonia serve as the bacterial reservoir in TB pneumonia. Acid-fast microscopy often detects few bacilli in patient biopsies^18^, raising the possibility that organisms persist in non-replicating or masked states. To address this, we employed in situ hybridization for *Mtb* mRNAs.

RNAscope for multiple *Mtb* transcripts (e.g., esxA, fbpB/85B) revealed clusters of mRNA signals within macrophages in the alveolar pneumonia, even in areas where acid-fast staining was negative (**Figure 5D; Supplemental Figure 14**)^46^. The pattern of *Mtb* mRNA staining was similar to the detection of PPD (**Figure 5C**). By contrast, in organized granulomas we observed only sparse *Mtb* mRNA puncta, typically as single dots in a few macrophages. Thus, the highest bacterial burden resided in the foamy macrophages in the alveolar pneumonia, not in cores of the granulomas. Taken together, the antigen staining and RNA hybridization provide compelling evidence that TREM2⁺ foamy macrophages in the alveolar pneumonia represents a host cell reservoir for *Mtb* in the lungs of pulmonary TB patients.

### What is the role of TREM2⁺ macrophages in TB pathogenesis?

Based on the spatial data showing TREM2⁺ foamy macrophages as a reservoir of *Mtb* mRNA and antigen in vivo, we next sought to determine how TREM2 is induced and whether its signaling is activated by pathogen-derived ligands. TREM2 is a receptor that senses lipids and apoptotic debris^47, 48, 49^. Two classes of signals were considered: (i) host cytokines, since IL-4/IL-13 drive alternative macrophage activation and upregulate TREM2^50, 51^, and (ii) *Mtb* cell-envelope lipids. In vitro, we confirmed that IL-4 treatment increased TREM2 expression on ∼30% of monocyte-derived macrophages, consistent with prior studies^50^. We then tested two major *Mtb* lipids: PDIM and free mycolic acids. Both potently induced TREM2 surface expression (∼50-55% of cells), a level surpassing even live *Mtb* infection (∼30% at MOI 5-10) **(Figure 6A and Supplemental Figure 17A)**. By contrast, lipoarabinomannan (LAM) and peptidoglycan failed to induce significant TREM2. Notably, PDIM induced TREM2 expression more strongly than live *Mtb*, identifying PDIM, a known virulence determinant, as a potent upstream driver of this foamy macrophage program. IL-4 induced TREM2 surface expression in ∼55-85% of cells (**Supplemental Figure 17A**), such that in future experiments we selected donors that reproducibly generated >80% TREM2 macrophages.

**Figure 6.**
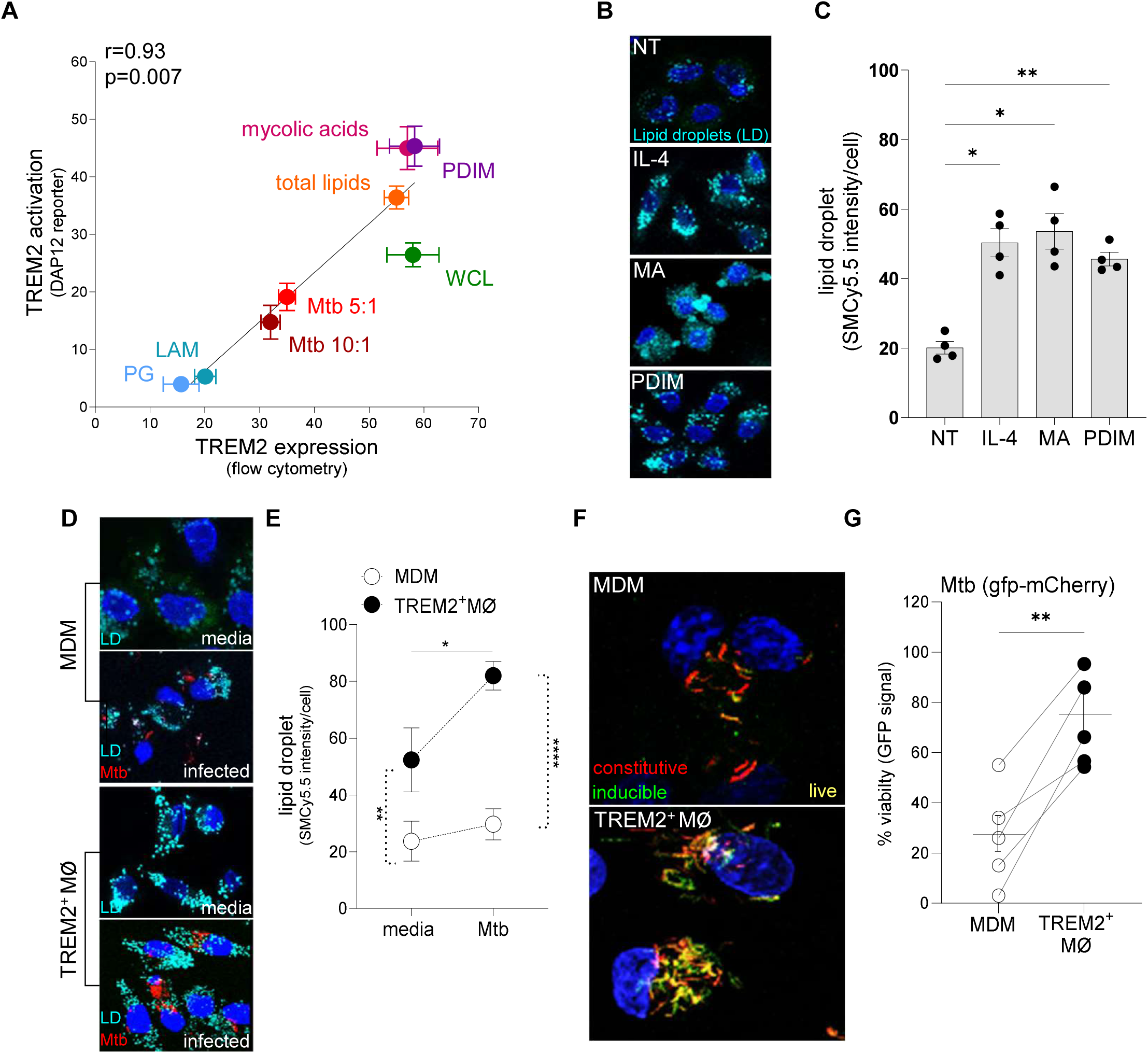
TREM2^+^ macrophage (MØ) interactions with *M. tuberculosis* (*Mtb*). (A) TREM2 activation correlates with its upregulation in monocyte-derived macrophages (MDMs) stimulated with *Mtb*-associated ligands (n = 7). (B-C) Mycolic acids (MA) and phthiocerol dimycocerosates (PDIM) induce lipid droplet (LD) aggregation at levels comparable to IL-4-polarized TREM2^+^ MØ. (D–E). Fluorescent images are representative of 5 individual donors. TREM2+ MØ exhibit a higher accumulation of lipid droplets than MDMs in uninfected and *Mtb* (mCherry)-infected conditions (n = 5). Lipid droplets (LD) were labeled with SMCy5.5 (cyan). The dotted lines connect the media and Mtb conditions of the difference macrophage populations, MDM and TREM2+ macrophages. (F–G) *Mtb* preferentially proliferates in TREM2+ MØ compared to MDMs. MØ were infected with *Mtb* (GFP-mCherry), with inducible GFP and constitutive mCherry expression (n = 5). The lines connect individual donors in which MDM and TREM2 macrophages were compared. Two-way ANOVA with Fisher multi-comparison test. Statistical significance: * p ≤ 0.05, ** p ≤ 0.01, ***p ≤ 0.001, ****p ≤ 0.0001.

We next assessed whether these lipids also activated TREM2 signaling. Using a fluorescent TREM2–DAP12 reporter, which clusters upon ITAM–SYK pathway engagement^23^, we found that both PDIM and free mycolic acids robustly triggered intracellular TREM2–DAP12 activation, with a strong correlation between surface TREM2 induction and signaling activity (r = 0.93, p = 0.007) (**Figure 6A and Supplemental Figure 17B)**. In contrast, LAM and peptidoglycan produced no significant reporter activation. Similarly, *Mtb* trehalose dimycolate (cord factor) did not induce TREM2 activation^22^. These results establish that specific *Mtb* lipids not only induce TREM2 expression but also directly stimulate its signaling cascade. These results demonstrate that PDIM and mycolic acids are not merely structural lipids but are active ligands that directly trigger TREM2–DAP12 signaling.

Because lipid droplets and the foamy state characterize TREM2⁺ macrophages in alveolar pneumonia, we next tested whether TREM2-inducing ligands also promote lipid storage. Infection of macrophages with *Mtb* triggers lipid droplet biogenesis^18^, consistent with immunometabolic reprogramming^52^. Using SMCy5.5, a lipid-sensitive fluorogenic probe that exhibits a 300–1,000-fold increase in fluorescence in lipid-rich environments, we observed an approximate 2.5-fold increase in lipid droplet formation after stimulation with IL-4, mycolic acids, or PDIM (**Figure 6B**). Oil Red O staining confirmed that each of these ligands significantly increased lipid droplet accumulation in human MDMs (**Supplemental Figure 18A–B**). Direct infection of TREM2⁺ macrophages with *Mtb* further increased lipid droplet abundance by ∼50% relative to uninfected TREM2⁺ macrophages **(Figure 6D–E**). Although mycolic acids were previously shown to induce lipid droplets^18^, these findings newly identify PDIM, a key virulence lipid^24, 53, 54, 55, 56, 57^, as a potent driver of the foamy TREM2⁺ macrophage phenotype.

### Do TREM2⁺ macrophages support or restrict *Mb* growth?

The spatial data indicated that TREM2⁺ foamy macrophages coincide with the highest bacillary burden in vivo. We considered why foamy, TREM2⁺-containing macrophages would favor the bacteria? One hypothesis is that excess lipid droplets and associated metabolic changes might impair bactericidal functions of macrophages while providing nutrients for *Mtb*. We observed in tissue that TREM2⁺ foamy macrophages co-localized with abundant *Mtb* antigens and mRNAs, whereas more classically activated macrophages in granulomas did not. To directly test the functional relationship between the foamy macrophage phenotype and *Mtb* viability, we conducted infection experiments in vitro.

Human macrophages were pre-differentiated with IL-4 to induce TREM2 expression and lipid accumulation (mimicking foamy macrophages in the alveolar pneumonia) or kept as unpolarized controls. We then infected both groups with a fluorescent *Mtb* reporter strain that expresses GFP preferentially in live (actively respiring) bacteria and mCherry constitutively^58, 59^. After 24 hours, confocal microscopy revealed that macrophages with high TREM2 and lipid content harbored approximately twice as many viable (GFP⁺) bacilli per cell compared to control macrophages (**Figure 6F, G**, **Supplemental Figure 19**). In other words, *Mtb* survival was significantly higher inside foamy TREM2⁺ macrophages. This result was consistent across multiple donors. We quantified the GFP/mCherry fluorescence ratio and observed a clear skew toward GFP (indicating live bacilli) in TREM2⁺ foamy macrophages. Thus, functional assays demonstrate that the TREM2⁺ foamy state directly increases intracellular *Mtb* viability.

We next asked whether immunometabolic reprogramming of the TREM2⁺ macrophages would enhance their bactericidal activity. Because we had previously shown that vitamin D can induce antimicrobial programs in *Mtb*-infected macrophages^33^, we asked whether it could reverse the permissive TREM2⁺ foamy phenotype. Remarkably, 1,25-dihydroxyvitamin D₃ (1,25(OH)₂D₃) pretreatment significantly downregulated TREM2 on macrophages (by flow cytometry) (**Supplemental Figure 20A, B**).

To determine whether this reprogramming improved bacterial control, we quantified intracellular survival by CFU assay. Vitamin D₃ pretreatment produced a robust antimicrobial effect, reducing bacterial burden by approximately 1 log₁₀ by day 4 in both TREM2⁺ macrophages and unsorted MDMs (p < 0.0001) (**Figure 7A**). We next used flow cytometry to define infection outcome and host cell viability within TREM2^+^ and TREM2⁻ macrophages (**Figure 7B**). After gating on live cells and stratifying into TREM2⁺ (TREM2⁺ MØ) and TREM2⁻ (MDM) populations, we found that *Mtb* infection resulted in a significantly higher fraction of TREM2⁺ macrophages harboring viable intracellular bacteria compared with TREM2⁻ cells. This indicates that enhanced bacterial persistence is an intrinsic feature of the TREM2-associated macrophage state rather than a consequence of differential cell death. Importantly, treatment with 1,25(OH)₂D₃ markedly shifted this distribution. Vitamin D reduced the frequency of infected TREM2⁺ cells while increasing the proportion of viable, bacteria-restricted macrophages, consistent with restoration of antimicrobial function. This shift occurred without a reduction in overall cell viability, demonstrating that vitamin D enhances bacterial control through functional reprogramming rather than cytotoxicity. Together, these data establish that the permissive TREM2⁺ macrophage state is functionally reversible and that vitamin D restores antimicrobial activity at both the single-cell and population levels.

**Figure 7.**
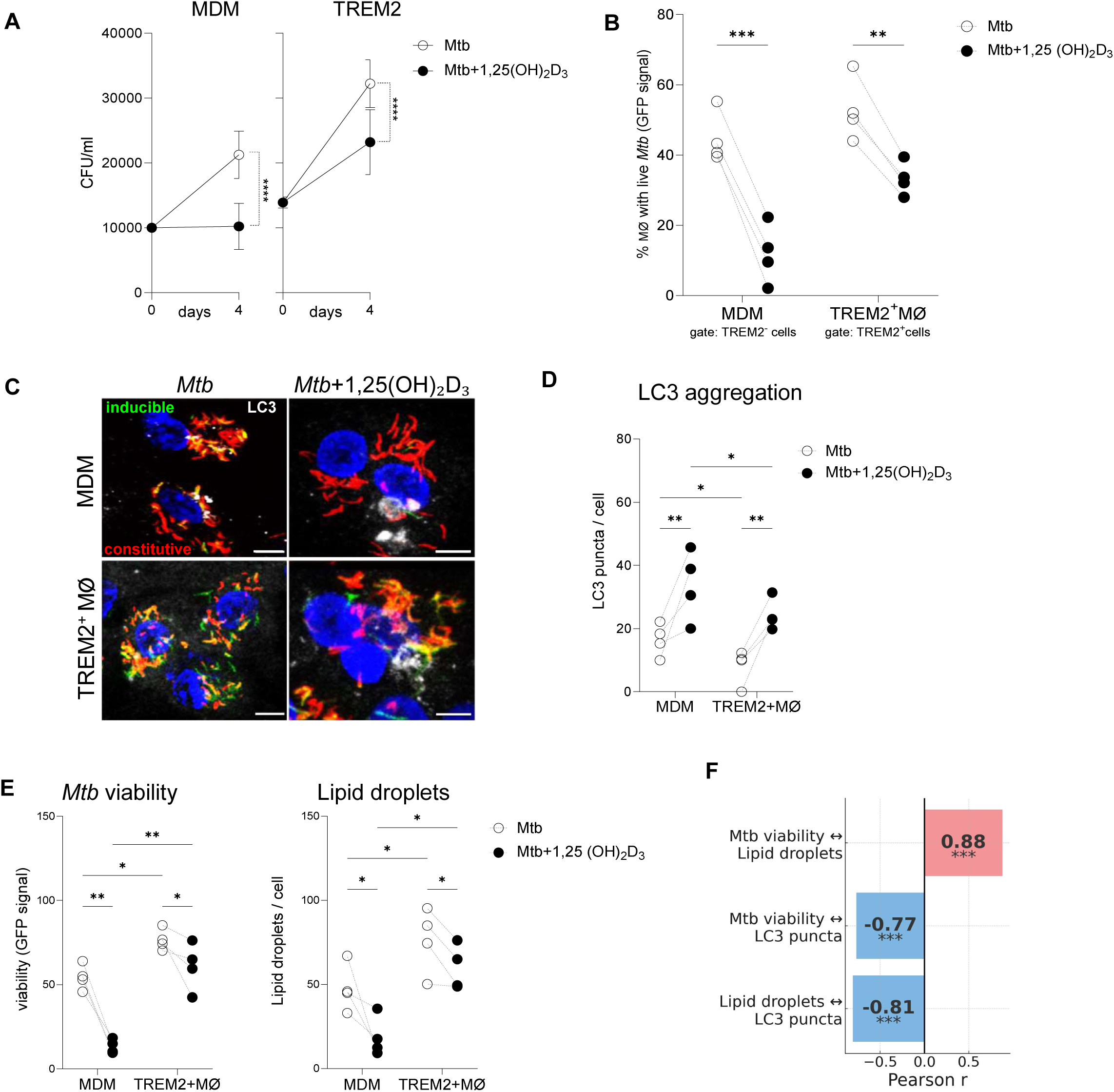
Vitamin D reduces *Mycobacterium tuberculosis* (*Mtb*) viability and lipid droplet accumulation by enhancing autophagy in TREM2⁺ macrophages: (A) antimicrobial activity of MDM and TREM2⁺ macrophages treated with 1,25-dihydroxyvitamin D₃ (1,25(OH)₂D₃) and infected with *Mtb* quantified by CFU assay (n = 5 donors); (B) flow cytometry analysis of donor-matched cells showing live-cell gating and quantification of infected TREM2⁺ and TREM2⁻ populations with or without vitamin D treatment; (C) representative confocal images of infected macrophages showing LC3 puncta around *Mtb* (white) and nuclei (DAPI); (D) quantification of LC3 puncta per cell from confocal images performed in ImageJ using experiments conducted side by side in the same donors (n = 4); (E) quantification of *Mtb* viability (GFP signal) and lipid droplets (SMCy5.5 staining) in MDM and TREM2⁺ macrophages infected with dual-reporter *Mtb* (GFP–mCherry), with lines indicating paired donor responses comparing *Mtb* alone versus 1,25(OH)₂D₃ plus *Mtb*; and (F) Pearson correlation analysis of donor-matched data (n = 16 points) demonstrating a positive association between lipid droplets and *Mtb* viability and inverse correlations between LC3 puncta and both lipid droplets and bacterial viability, with statistical analysis performed using two-way ANOVA with Fisher’s multiple-comparison test (p ≤ 0.05, p ≤ 0.01, p ≤ 0.001).

To define the mechanism underlying the effect of vitamin D, we examined LC3 recruitment during infection. Vitamin D treatment induced a marked increase in LC3 puncta formation in infected macrophages, indicating activation of an autophagy-dependent antimicrobial program (**Figure 7C**). Vitamin D has been shown to promote ULK1- and Beclin-1-dependent canonical autophagy in human macrophages through induction of the antimicrobial peptide cathelicidin (LL-37), a pathway required for control of *Mtb*^60, 61, 62^. Consistent with this mechanism, increased LC3 recruitment was accompanied by enhanced bacterial restriction, supporting canonical autophagy as a central pathway through which vitamin D restores antimicrobial function in TREM2⁺ macrophages. Quantitative LC3 analysis confirmed that TREM2⁺ macrophages maintain lower basal autophagy than MDMs, while 1,25(OH)₂D₃ increased LC3 puncta in both populations. Thus, *Mtb*-infected TREM2⁺ macrophages contain more viable bacilli, accumulate more lipid droplets, and display reduced autophagy relative to MDMs (**Figure 7D**), features that 1,25(OH)₂D₃ effectively reverses.

Metabolic characterization performed side by side with LC3 analyses in the same donor cohort, revealed that TREM2⁺ macrophages accumulated markedly increased lipid droplets compared with unpolarized monocyte-derived macrophages (MDMs). TREM2⁺ macrophages exhibited a 2–3-fold increase in lipid droplet content (SMCy5.5 staining; p < 0.05; n = 4 donors), consistent with the foamy metabolic state observed in vivo. In parallel *Mtb* viability reporter assays conducted under the same experimental conditions (**Figure 7E and Supplementary Fig. 21**), this lipid-rich phenotype was associated with 30–50% higher GFP/mCherry ratios, indicating a greater proportion of metabolically active intracellular bacilli. Elevated lipid content therefore strongly correlated with increased bacterial viability, supporting the concept that lipid accumulation creates a metabolically favorable intracellular environment for *Mtb* persistence. Treatment with 1,25(OH)₂D₃ reprogrammed both macrophage populations, reducing lipid droplet abundance by ∼40–55% (p < 0.05) and concomitantly decreasing GFP/mCherry viability ratios by 30–50% (p < 0.01), consistent with coordinated metabolic and antimicrobial restoration.

To further assess the relationships among lipid metabolism, autophagy, and intracellular *Mtb* survival, we performed correlation analyses across donor-matched datasets. Lipid droplet abundance correlated strongly and positively with *Mtb* viability (**Supplementary Fig. 22**; r = 0.86, p = 0.0002), indicating that cells with greater lipid stores harbor a higher proportion of metabolically active bacilli. In contrast, LC3 puncta showed a robust inverse correlation with both lipid droplet content (r = –0.81, p = 0.0008) and *Mtb* viability (r = –0.77, p = 0.0009), consistent with the antagonism between autophagy and lipid accumulation. These coordinated relationships, *Mtb* viability correlating with lipid droplets and inversely with autophagy, reinforce the linkage between the TREM2⁺ foamy state and enhanced bacterial fitness (**Figure 7F**).

In summary, our functional experiments demonstrate that the TREM2⁺ foamy macrophage state directly contributes to *Mtb* survival in alveoli (**Figure 8**). *Mtb*’s own lipids increase *Mtb* survival in macrophages; an intervention that counters this (such as vitamin D) restore macrophage ability to suppress the bacteria. Together with the spatial data, these results paint a coherent picture of what may occur in early TB infection, *Mtb* co-opts macrophages in the alveolar pneumonia to become foamy lipid-loaded cells (via TREM2 and other pathways), which then serve as incubators for the bacteria, all while mounting a muted immune response. This foamy macrophage stage likely persists as the dominant bacillary niche until the host’s adaptive response is engaged and granulomas form.

**Figure 8.**
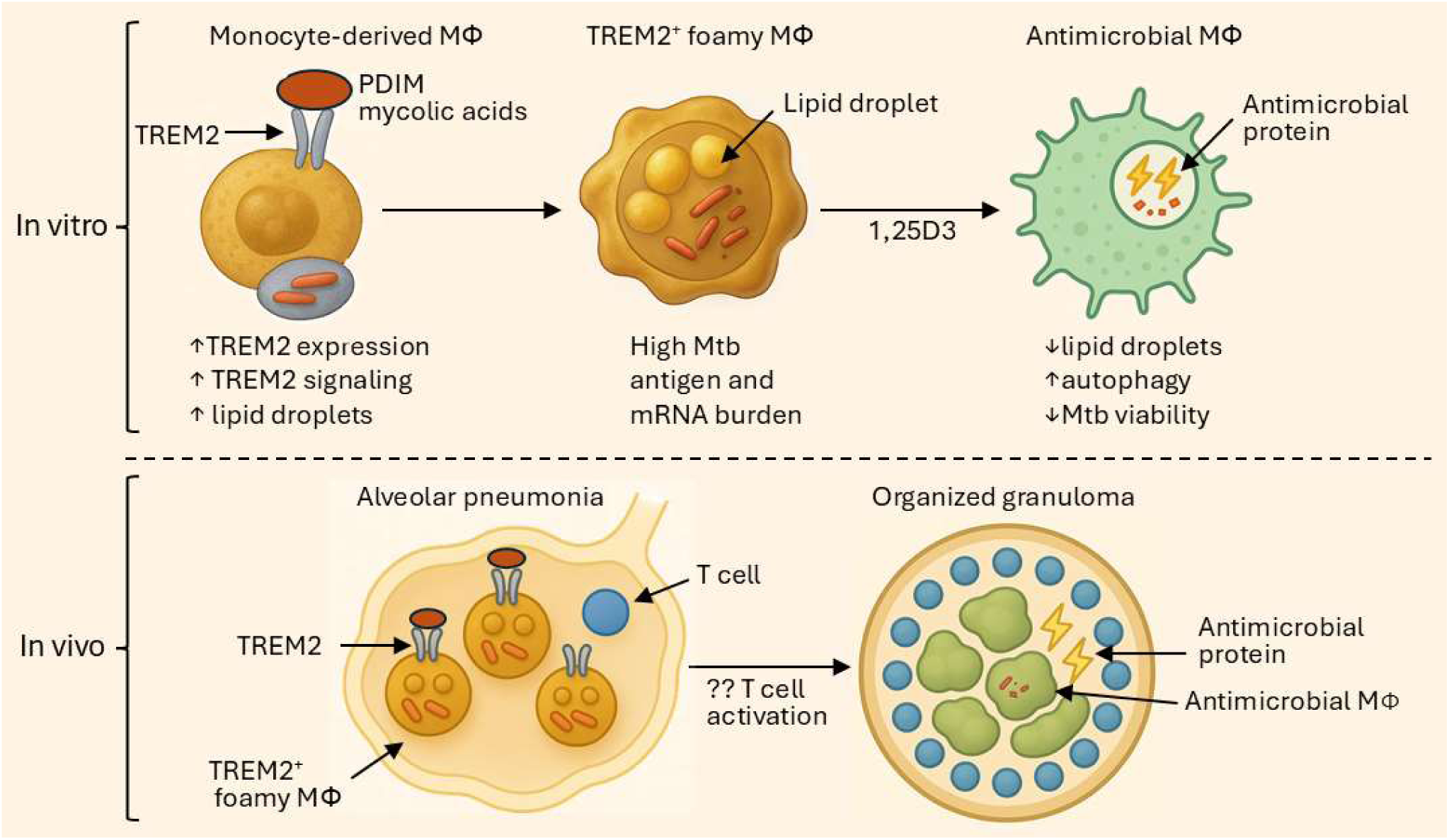
Reprogramming of TREM2 foamy macrophages into antimicrobial macrophages in tuberculosis. PDIM-induced TREM2 signaling drives formation of lipid-rich foamy macrophages that harbor a high Mtb burden. Treatment with 1,25(OH)₂D₃ reprograms these cells into antimicrobial macrophages with increased autophagy and reduced Mtb viability. In vivo, TREM2⁺ foamy macrophages dominate early alveolar pneumonia and precede the emergence of granuloma-associated antimicrobial macrophages including IFN-γ-activated macrophages and SPP1^+^ macrophages.

## Discussion

Spatial transcriptomics of human pulmonary tuberculosis biopsy specimens established that alveolar pneumonia and granuloma cores represent fundamentally divergent immunologic states within the same lung. Alveolar regions were enriched for lipid metabolism and scavenger receptor programs, including APOE, APOC1, CD163, and LIPA, consistent with TREM2-associated macrophage states described in lipid-rich pathologies^30, 63^. The alveoli in TB pneumonia were largely devoid of T cells, a notable finding, as it indicates bacillary expansion occurs in a compartment with minimal adaptive immune pressure. By contrast, granuloma cores showed robust induction of hallmark antimicrobial pathways (IL1B, CAMP, CYP27B1, CYBB) and enrichment of IFN-γ–activated macrophages and Th17 cells. Notably, transcripts encoding antimicrobial peptides were relatively absent from alveoli but abundant in granulomas. This dichotomy was visually confirmed by immunohistology and CosMx spatial proteomics: CD163⁺, PLIN2⁺, TREM2⁺ foamy macrophages densely occupied alveolar spaces, while granulomas contained epithelioid macrophages, multinucleated giant cells, and organized T and B cell mantles. This transcriptional observation was complemented by functional assays: In vitro, macrophages enriched in the TREM2⁺ foamy phenotype harbored approximately many more viable bacilli as unpolarized controls. Together, these findings indicate that TREM2⁺ macrophages form a permissive niche that enables persistence and early transmission of *Mtb*.

A central advance of this study is the demonstration that the *Mtb* virulence lipid PDIM, and, as previously shown, free mycolic acids^22, 23^, are potent inducers of TREM2 expression, lipid droplet biogenesis, and TREM2–DAP12 signaling in human macrophages, in some contexts exceeding the effects of live *Mtb* infection. These findings suggest a positive feedback loop in which cell envelope lipids engage TREM2 signaling, further amplifying TREM2 expression and suppressing macrophage antimicrobial function, consistent with a prior report^23^ and our data. PDIM is already recognized as a multifunctional virulence lipid that traffics into host membranes, inhibits phagocytic clearance, and cooperates with ESX-1 to disrupt phagosomal integrity^43, 55, 57, 64^. Our findings extend this paradigm by identifying TREM2 induction and lipid metabolic reprogramming as additional PDIM-associated effects that reshape macrophage identity. Together, these data reinforce PDIM as a pleiotropic virulence factor coordinating multiple host-directed pathways to promote macrophage states permissive for *Mtb* persistence.

Mechanistically, PDIM and mycolic acids triggered TREM2-dependent remodeling through a DAP12–PI3K–AKT–mTOR axis, promoting lipid uptake, lipid droplet accumulation, and suppression of autophagy, a program reminiscent of TREM2⁺ macrophage states described in neurodegeneration and metabolic disease^65, 66^. Consistent with spatial profiling, in vitro TREM2⁺ macrophages accumulated markedly more lipid droplets and exhibited reduced autophagic activity compared with unpolarized controls. This metabolic state directly favored intracellular survival: lipid droplet abundance strongly correlated with bacterial viability across donors (r = 0.98), and TREM2⁺ foamy macrophages harbored nearly twice as many viable bacilli as controls. These findings align with prior studies demonstrating that TREM2 signaling suppresses antimicrobial activity in mouse^22^ and human^23^ macrophages, collectively supporting a functional role for TREM2 in establishing a permissive intracellular niche.

Importantly, this pathogenic state was not irreversible: We show it is pharmacologically reversible by in vitro treatment with 1,25(OH)₂D₃ which downregulated TREM2 surface expression, reduced lipid droplets by ∼40–55%, increased autophagy, and reduced intracellular *Mtb* viability by ∼1 log_10_ CFU. 1,25(OH)₂D₃ did not fully restore the antimicrobial activity, consistent with the ability of *Mtb* to induce type I IFN in TREM2 macrophages^23^, a known inhibitor of the vitamin D antimicrobial pathway^67^. Nevertheless, to date, the clinical studies of vitamin D as a prevention or treatment of tuberculosis has yielded disappointing results^68, 69, 70, 71,72^. Our data highlight that the TREM2 macrophage permissive niche in human TB is modifiable and therefore represents a viable target for host-directed therapies.

In contrast to alveolar pneumonia, the granuloma exhibited marked upregulation of antimicrobial genes, defining a spatially organized immune niche resistant to *Mtb* growth. This architecture is consistent with spatial proteomic analyses demonstrating concentration of antimicrobial effector proteins within human TB granulomas^15^. Bulk transcriptomic profiling of resected human TB lesions similarly identified interferon/cytokine–dominated inflammatory modules associated with disease severity^73^, an immune program that aligns closely with the IFN-γ–activated macrophage and Th17 networks we localize to organized granuloma cores. In addition to canonical IFN-γ–mediated activation, CD4 T cells can promote macrophage antimicrobial function through GM-CSF–dependent, HIF-1α–mediated metabolic reprogramming, underscoring the importance of T cell–driven macrophage metabolic activation in controlling *Mtb* growth^74^. Likewise, single-cell profiling of bronchoalveolar lavage cells from *Mtb* “resisters” revealed coordinated M1 macrophage and poly-cytotoxic IFN-γ–producing T cell responses^75^, reinforcing the concept that Th1-polarized macrophages and cytotoxic lymphocytes define antimicrobial containment programs in human TB. Spatially resolved human granuloma atlases further substantiate the presence of Th1/Th17-enriched immune microenvironments within TB lesions^14, 15, 45^.

The patients studied here had negative sputum smears and cultures, typical of early or atypical TB presentations and biopsies were required to establish diagnosis. Despite the absence of acid-fast bacilli on routine staining, biopsy specimens of these lesions contained abundant *Mtb* transcripts and antigens. The diagnostic weight of mRNA is particularly strong, as transcripts are highly labile (genome-wide half-life ∼9.5 minutes in vitro) and clear rapidly once bacilli lose viability^76, 77^; clinical studies show mRNA disappears early during therapy, before DNA levels fall^78^. Antigen detection in tissue, even when AFB stains are negative, further supports the presence of metabolically active organisms^46^. Recent longitudinal cohorts demonstrate that many asymptomatic, culture-negative individuals harbor *Mtb* biomarkers, including mRNA transcripts and growth in augmented cultures, despite negative standard cultures^79^. Importantly, these organisms were found almost exclusively in the alveolar pneumonia, not in granuloma cores. Our findings thus provide a mechanistic explanation for how bacilli can expand, persist, and be exhaled before classical granulomas form and before symptoms arise: the alveolar niche dominated by TREM2⁺ foamy macrophages is permissive, metabolically supportive, and spatially positioned at the air–lung interface. The concept that lung-specific microenvironmental remodeling, rather than systemic immune failure, governs post-primary progression is further supported by experimental models demonstrating that dysplastic epithelial repair and maladaptive macrophage polarization uniquely predispose lung tissue to necrotizing TB^80^.

TB remains a leading cause of infectious mortality, in part because transmission frequently occurs from individuals without overt symptoms. Community-based surveys indicate that up to 80% of bacteriologically confirmed pulmonary TB among exposed adults may be asymptomatic^81, 82^, and viable bacilli can be released during normal tidal breathing^83^. Consistent with these observations, clinic- and community-based surveys in South Africa demonstrated that the majority of individuals with culture-positive sputum reported no TB symptoms^84^. Classical pathology established that the earliest pulmonary lesion is an exudative alveolar process preceding organized granuloma formation^6, 7, 9^. However, the cellular and molecular architecture of this early pneumonia has remained incompletely defined.

## Methods

### Patient selection and study approval

We collected pulmonary tissue specimens from patients at Charles Nicolle Hospital (Tunis) who presented with suspected tuberculosis (including atypical, acid-fast bacilli–negative, pseudotumor cases). Patients were enrolled at the time of diagnosis, before initiating therapy. The study was conducted in accordance with the Declaration of Helsinki and approved by the Institut Pasteur de Tunis Institutional Review Board (2021/26/I/V1) and the Charles Nicolle Hospital Institutional Review Board (2022/06/EC/V1). All participants provided written informed consent for blood and tissue collection.

### FFPE tissue preparation

We processed tissue samples immediately after surgical excision. Specimens (approximately 4–5 mm thick) were fixed in 10% neutral-buffered formalin (NBF) at room temperature for 24 h to preserve morphology. Fixed tissues were dehydrated through a graded ethanol series (70% ethanol for 1 h; 95% ethanol for 2 × 1 h; 100% ethanol for 2 × 1 h) and cleared in xylene. We then infiltrated tissues with molten paraffin wax at 58–60°C for 1 h under gentle vacuum. Samples were embedded in metal molds with paraffin, oriented for proper sectioning, and allowed to solidify at room temperature. Paraffin blocks were sectioned at 4–5 µm thickness on a rotary microtome. Sections were floated on a 42°C water bath, mounted onto positively charged glass slides, and dried (overnight at 37°C or 1–2 h at 60°C) to ensure adherence.

### GeoMx Digital Spatial Profiler (DSP) RNA

Formalin-fixed, paraffin-embedded (FFPE) tissue sections (5 µm) were mounted on glass slides for spatial transcriptomic analysis using the GeoMx Digital Spatial Profiler (DSP) platform (NanoString). Sections were deparaffinized and rehydrated according to the manufacturer’s protocols. For spatial RNA profiling of specific regions of interest (ROIs), we hybridized sections overnight at 37 °C with the GeoMx Human Whole Transcriptome Atlas (WTA) probe set (NanoString Technologies), which contains > 18,000 barcoded RNA probes targeting human protein-coding genes. Hybridization was carried out in a specialized buffer to promote specific RNA–probe binding and minimize nonspecific interactions.

After hybridization, tissue morphology and cellular markers were revealed by fluorescent immunostaining, and high-resolution fluorescence images were acquired on the GeoMx DSP. ROIs were selected based on these images, and targeted in situ transcript counting was performed: UV light was used to photocleave and release the barcoded probes bound to mRNA in each ROI. The released oligonucleotide probes from each ROI were collected into separate wells of a microtiter plate. For morphological context, tissue sections were stained with fluorescent markers (DAPI and immune cell markers) to guide ROI selection. In sample PTB 21.1, ROIs contained 400–1000 nuclei and were stained with DAPI, CD68, CD3, and CD8; in samples PTB 22.2, PTB 22.1, and PTB 22.3, ROIs contained ∼100 nuclei and were stained with DAPI, CD68, CD3, and CD20 (CD20 in place of CD8). Spatial transcriptomic libraries were generated from ROI eluates. Paired-end reads were produced (Illumina sequencing) and aligned and quantified using NanoString’s DSP analysis pipeline. Transcript counts were output in digital count conversion (DCC) format, linking reads to target genes.

### GeoMx data analysis

Raw DSP read counts (DCC files) were processed in R using GeomxTools (v3.10.0), GeoMxWorkflows (v1.12.0), and NanoStringNCTools (v1.14.0) with the GeoMx WTA probe annotations (v1.0). For initial processing, we pooled our pulmonary tuberculosis (PTB) samples with additional tuberculosis lymphadenitis and extrapulmonary TB samples to improve normalization; the extra samples were excluded from final analysis after quality control (QC) filtering. We applied default NanoString QC criteria for ROIs: each ROI required ≥ 1,000 sequencing reads, ≥ 80% probe reads passing trimming and stitching, ≥ 75% reads aligning to targets, ≥ 1 positive control count, ≤ 1,000 counts in the no-template control well, ≥ 20 nuclei per ROI, and ROI area ≥ 1,000 µm. One ROI (core region) failed these criteria and was removed. We defined the per-ROI background level as the limit of quantification (LOQ), calculated as the geometric mean of negative control probe counts multiplied by the geometric variance. Anyone-inflated gene counts below the LOQ (or below a minimum threshold of 2) were considered undetectable. We filtered out ROIs that did not have at least 5% of all WTA genes above background expression (8 ROIs were removed for low gene detection). Genes not detected in at least 5% of the remaining ROIs were also removed. Starting from 18,677 targeted genes in the WTA, 10,968 genes passed these filtering steps.

As our analysis focused on comparing alveolar, core, and mantle regions within PTB granulomas, we excluded ROIs that were not clearly from one of these three regions. One ROI initially classified as “core” was excluded due to partial contamination with mantle tissue. ROIs identified as “infiltrate” or “fibrotic mantle” were retained for modeling but not included in direct alveoli/core/mantle comparisons. In total, 79 ROIs were profiled from PTB lesions; after QC filtering, 70 ROIs remained, of which 59 were confidently assigned as alveoli (18 ROIs), core (25 ROIs), or mantle (14 ROIs).

### GeoMx gene signatures

Differential gene expression analysis was performed to identify region-specific transcriptional signatures. We used the DESeq2 R package (v1.46.0) to model gene counts with an additive linear model including region (alveoli, core, mantle) and patient (PTB 21.1, 22.1, 22.2, 22.3) as factors. From this model, we extracted lists of differentially expressed genes (DEGs) for each pairwise region comparison (adjusted P ≤ 0.05). We defined each region’s signature gene set as the genes upregulated in that region compared to both of the other regions. For example, the “alveoli” gene set was the intersection of DEGs from alveoli vs core and alveoli vs mantle comparisons.

We performed dimensionality reduction to visualize ROI-level transcriptomic relationships. We applied a variance-stabilizing transformation (VST) to the normalized counts and selected the 500 most variable genes across all alveoli, core, and mantle ROIs. Principal component analysis (PCA) was conducted on these genes, and the top principal components were used to generate a two-dimensional Uniform Manifold Approximation and Projection (UMAP) embedding. This UMAP visualization highlights the segregation of ROIs by region based on their gene expression profiles.

### scRNA -seq from PTB FFPE

We performed single-cell RNA sequencing (scRNA-seq) on four FFPE pulmonary tuberculosis tissue samples (PTB 21.1, PTB 22.1, PTB 22.2, PTB 22.3). From each formalin-fixed specimen, 15 µm scrolls were sectioned and processed following the 10x Genomics Fixed RNA Profiling sample preparation protocol. In brief, deparaffinized tissue scrolls were mechanically dissociated with a tissue grinder (pestle) and enzymatically digested with Liberase to release nuclei. A single-cell/nucleus suspension was obtained for each sample.

Single-cell libraries were prepared using the 10x Genomics Chromium Fixed RNA Profiling (GEM-X) chemistry. Prior to probe hybridization, each sample yielded on average ∼3.5×10^5 nuclei; after probe hybridization and washes, an average of ∼1.9×10^5 nuclei per sample remained. Equal numbers of nuclei from each sample were then combined to create a multiplexed library, ensuring roughly 25% representation from each of the four patient samples. Unique oligonucleotide barcodes were assigned to each sample to enable demultiplexing. Libraries were sequenced on an Illumina NovaSeq X Plus platform (paired-end 2 × 50 bp reads), generating approximately 1.25 billion total reads for the pooled library.

### scRNA-seq from PTB FFPE analysis

Raw sequencing data were processed with the 10x Genomics Cell Ranger pipeline. Four samples (PTB 21.1a, PTB 21.1b, PTB 22.2, PTB 22.3) were demultiplexed, aligned, and gene-count matrices generated using Cell Ranger v9.0.1 against the GRCh38-2024-A human reference. The remaining sample (PTB 22.1, which had been sequenced in a separate run) was processed with Cell Ranger v7.2.0 against the GRCh38-2020-A reference. The count matrices were imported into the Seurat R package (v5.1.0) for quality control and downstream analysis. We corrected for ambient RNA contamination using SoupX (default parameters) before filtering. Low-quality cells/nuclei were removed by requiring each barcode to have ≥ 400 detected genes and ≤ 5% mitochondrial RNA content. The filtered datasets from all samples were then merged. Data normalization and scaling were performed in Seurat, and the top variable genes were used for principal component analysis. We integrated the samples to account for batch effects using the Harmony algorithm (Harmony integration with default settings). Significant principal components were used to construct a shared nearest-neighbor graph, and we obtained two-dimensional embeddings with Uniform Manifold Approximation and Projection (UMAP). Clustering was performed using Seurat’s graph-based clustering (FindNeighbors and FindClusters) at a low resolution (0.1), and cluster robustness was evaluated with the Clustree tool. We assigned cluster identities by examining canonical cell-type marker genes and confirmed these via differential expression testing (Seurat FindAllMarkers).

We next extracted major immune cell types of interest for sub-clustering. In particular, we separately removed the macrophage/monocyte lineage and the T cell lineage from the integrated dataset and re-ran the normalization, PCA, Harmony integration, and clustering workflow on each subset individually (maintaining the same filtering thresholds). Clustering resolution was optimized (guided by Clustree) and set to 0.3 for both macrophages and T cells to identify fine-grained subpopulations. Subclusters were annotated based on their top differentially expressed genes (FindAllMarkers results). Finally, these annotated subclusters were mapped back to the full dataset for comparative analysis with the spatial transcriptomic data.

### Jaccard overlap of GeoMx and scRNA-seq data

To assess biological concordance between spatially defined transcriptional programs and cell type–specific expression signatures, we quantified the overlap between GeoMx-derived granuloma region signatures (alveoli, core, and mantle) and curated gene sets from single-cell RNA-seq (scRNA-seq) datasets. These included scRNA-seq signatures from pulmonary tuberculosis lesions, human leprosy granulomas containing TREM2⁺ macrophages^29^, a healthy human lung cell atlas^28^, and published TREM2⁺ macrophage–associated gene signatures.

For each region-specific DEG set, we computed Jaccard indices (intersection over union) relative to each reference signature to quantify transcriptional similarity. Global patterns of overlap were visualized by hierarchical clustering of the reference gene sets based on their Jaccard similarity to the alveolar, core, and mantle signatures (using average linkage and Spearman correlation as the distance metric). Statistical significance of gene set overlaps was assessed by hypergeometric testing, with multiple hypothesis correction using the Benjamini–Hochberg procedure.

Together, these analyses linked spatial gene expression profiles to known immune cell phenotypes, revealing conserved and regionally distinct programs. We observed enriched overlap with signatures of lipid-laden, TREM2⁺ macrophages and antimicrobial effector pathways highlighting their contribution to the spatial organization and functional compartmentalization of TB granulomas.

### Analysis of genes encoding antimicrobial peptides

We assembled a targeted list of genes encoding antimicrobial peptides/proteins to interrogate their expression patterns in PTB lesions. An initial list of 52 antimicrobial genes was compiled from the Antimicrobial Peptide Database (APD3)3 and literature on host defense in mycobacterial infection^28, 29, 31, 32^. Of these 52 genes, 48 were detected in either our PTB scRNA-seq dataset or the GeoMx spatial transcriptomic dataset (or both). We included four additional immune genes (CCL26, CCL27, S100A7, S100A7A) due to their known roles in mycobacterial infections, bringing the total list to 56 genes.

To explore relevant literature, we performed a comprehensive PubMed search for each antimicrobial gene in the context of tuberculosis. We developed a Python script that queried the NCBI E-utilities with the search string “tuberculosis AND (antimicrobial OR kill) AND (GeneName OR ProteinName)” for each gene. For each hit, the script retrieved the PubMed ID, title, authors, journal, year, abstract, and DOI. Results were compiled into a Pandas dataframe, duplicate entries were removed, and the final list was exported to a spreadsheet with live DOI links for reference.

We evaluated the overlap between region-specific DEGs in our spatial data and predefined antimicrobial gene sets. For each PTB region (alveoli, core, mantle), we computed the Jaccard index comparing the region’s DEG set to the list of antimicrobial genes. We also compared the antimicrobial gene list to the marker genes of key cell populations identified in the PTB scRNA-seq dataset (e.g., TREM2^+^ macrophages). The significance of overlap between gene sets was tested using hypergeometric tests. For all these analyses, gene expression counts were normalized with DESeq2 and transformed with VST as described earlier, to ensure comparability in gene selection and ranking.

### Functional gene analysis

We performed functional enrichment analyses to identify biological pathways and processes associated with differentially expressed genes in PTB lesions. The upregulated DEGs from two sources (Zhou, Y. et al 2019) – GeoMx spatial profiling and scRNA-seq – were analyzed separately using Metascape (v3.5) 5, a web-based gene annotation and enrichment tool. Each input gene list was filtered for significance (adjusted p < 0.05 and |log_2 fold-change| ≥ 1) prior to analysis. Metascape integrated multiple ontology sources (Gene Ontology, Reactome, WikiPathways, etc.) to compute overrepresentation of functional terms. We used default Metascape parameters, including a minimum of 3 genes per term, minimum enrichment score of 1.5, and a p-value cutoff of 0.01 for term inclusion.

Enriched terms were ranked by statistical significance and coverage, and redundant terms were clustered based on Kappa similarity scores. We visualized the top enrichment results as bar plots highlighting key biological processes and pathways for each dataset. Notably, both spatial and single-cell data revealed enrichment of immune activation pathways, including antigen presentation, cytokine signaling, antimicrobial responses, and tissue remodeling processes in PTB granulomas.

### TREM2 macrophage signature analysis in pulmonary TB spatial transcriptomic data

To assess representation of TREM2-associated macrophage programs in our spatial transcriptomic data, we compared our DEGs with published TREM2^+^ macrophage gene signatures. We curated 34 gene sets representing TREM2^+^ macrophage signatures from 21 independent studies. For each gene in these signatures, we calculated two metrics: its frequency of occurrence across the 34 signature gene sets, and its Jaccard index overlap with the DEGs from our GeoMx data (specifically the combined set of DEGs distinguishing alveolar vs core regions, which showed the strongest macrophage polarization differences).

We considered genes that appeared in ≥ 22 of the 34 literature signatures (approximately two-thirds of the signatures) to be highly recurrent features of TREM2^+^ macrophages. We examined whether these high-recurrence genes were differentially expressed in our alveoli vs core comparison. For each such gene, we calculated an expression ratio between alveolar and core regions using DESeq2-normalized counts (log10 of alveoli/core). Genes not detected in the GeoMx dataset were excluded from ratio calculations. This analysis allowed us to identify which canonical TREM2^+^ macrophage genes were enriched in different granuloma regions and to place our findings in the context of reported macrophage activation states.

### CosMx Spatial Molecular Imager (SMI) protein assay

We conducted high-plex spatial proteomic analysis on FFPE tissue sections using the NanoString CosMx Spatial Molecular Imager (SMI) platform. FFPE tissue sections (5 µm) mounted on glass slides were deparaffinized, rehydrated, and subjected to heat-mediated antigen retrieval according to the CosMx manufacturer’s protocol. We then incubated the sections with the CosMx Human Immuno-Oncology Protein Panel, which targets 64 proteins, by applying the cocktail of DNA-barcoded antibodies to each slide. Hybridization was carried out at 4 °C in a humidified chamber for 16–18 h. After incubation, slides were washed to remove unbound probes and reduce background fluorescence.

The slides were loaded onto the CosMx SMI instrument for automated imaging and data collection. An initial fluorescence scan (CosMx configuration A, pre-bleach settings) was performed to establish signal profiles, followed by iterative imaging cycles (using the instrument’s human (non-neural) segmentation configuration) to capture barcoded probe signals for each protein target at subcellular resolution. Tissue architecture and protein localization were visualized by counterstaining nuclei with DAPI and using a subset of fluorescently labeled morphology markers included in the panel. We selected multiple fields of view (FOVs) per sample based on histological features and regions of interest (e.g., caseating necrosis core, cellular rim).

The CosMx SMI system simultaneously quantified protein expression at the subcellular level by detecting the unique oligonucleotide barcodes attached to each antibody. After imaging, we used the CosMx SMI analysis pipeline to process the data. This pipeline segmented each FOV into subcellular compartments (nucleus, cytoplasm, membrane) using integrated image analysis algorithms, enabling quantification of protein expression within each cell compartment. Raw barcode counts for each protein were normalized to internal controls on a per-sample basis. The processed data were used to map and visualize the spatial distribution of all 64 proteins across the tissue. We also utilized NanoString’s AtoMx software to assist with cell segmentation and to interactively explore the spatial expression patterns of proteins of interest.

### Reagents

Cytokines for cell culture: Human recombinant IFN-γ (BD Biosciences, cat. no. 554616), M-CSF (R&D Systems, cat. no. 216-MC-100/CF), and IL-4 (Thermo Fisher Scientific, cat. no. 200-04-1MG) were used in macrophage differentiation and activation assays.

Flow cytometry antibodies: APC-conjugated anti-TREM2 monoclonal antibody (rat IgG2b isotype, R&D Systems, cat. no. FAB17291A) with its matched isotype control; PerCP-eFluor 710 (PerCP-A710)–conjugated anti-CD163 monoclonal antibody (mouse IgG1, Thermo Fisher, cat. no. 46-1639-42) with matched isotype control. Density gradient medium: Ficoll-Paque PLUS (GE Healthcare) was used for PBMC isolation. Primary antibodies for immunohistochemistry (IHC) and immunofluorescence: CD3 (rat IgG1, Bio-Rad, cat. no. MCA1477); CD68 (mouse IgG1, Thermo Fisher, cat. no. 14-0688-82); CD20 (mouse IgG2a, Thermo Fisher, cat. no. 14-0202-37); PLIN2/ADRP (mouse IgG1, Thermo Fisher, cat. no. MA5-24797); TREM2 (mouse IgG3, Novus Biologicals, cat. no. NBP1-07101); APOE (mouse IgG1, Novus, cat. no. NB110-60531); SPP1/osteopontin (mouse IgG1, Novus, cat. no. NB110-89062); Alexa Fluor 488–conjugated anti-TREM2 (directly labeled mAb, Novus, cat. no. NBP1-07101AF488); CD4 (rabbit IgG, Abcam, cat. no. AB133616); CD8 (mouse IgG1, Agilent Dako, cat. no. M710301-2); CD163 (mouse IgG2a, Lifespan Biosciences, cat. no. LS-B12939-50); *Mtb* lipoarabinomannan (LAM) antibody (mouse IgG3, BEI Resources, cat. no. NR-13811); polyclonal *Mtb* antibody (rabbit IgG, Biocare Medical, cat. no. CP140A) and LC3 (mIgG1) from MBL International (Woburn, MA, US).

Secondary detection reagents: Unconjugated IgG isotype control antibodies (Sigma-Aldrich); biotinylated anti-mouse IgG and anti-rabbit IgG secondary antibodies (Vector Laboratories, included in VECTASTAIN Elite ABC kits); normal horse serum and normal goat serum blocking reagents (Vector); VECTASTAIN Elite ABC peroxidase kit (Vector); AEC peroxidase substrate kit (Vector); Alexa Fluor 488, 568, and 647 goat anti-mouse IgG subclass-specific antibodies, and goat anti-rabbit IgG (Thermo Fisher) for immunofluorescence.

Cell isolation reagents: Anti-CD14 magnetic microbeads (Miltenyi Biotec) for monocyte isolation from PBMCs.

### RNA in situ hybridization

RNA in situ hybridization was performed to detect *Mtb* RNA in FFPE lung sections^46^. We used the RNAscope 2.5 HD Red kit (Advanced Cell Diagnostics, cat. no. 322372) following the manufacturer’s instructions, with a specific probe set for the RNAscope® Probe -B-*M.tuberculosis*-Erdman (Cat no. 552911), with Hs PPIB (PN 313901) as the positive control probe and dapB (PN 310043) as the negative control probe. FFPE tissue sections were deparaffinized and underwent target retrieval (boiling in retrieval buffer) for 30 min. Hybridization with the *Mtb* probe and control probes was carried out at 40°C in the HybEZ oven, followed by a series of signal amplification steps. We implemented a modified AMP 5 incubation of 45 min (extended from the standard protocol) to enhance signal intensity. After completing the RNAscope amplification and development steps (Fast Red chromogenic detection), slides were counterstained with 50% Gill’s hematoxylin I, then rinsed in water. Stained slides were mounted with EcoMount medium (Biocare Medical, cat. no. EM897L) and scanned at 80× magnification using a Motic EasyScan One digital slide scanner.

### Tissue immunoperoxidase labeling

We carried out immunoperoxidase (chromogenic IHC) staining on FFPE tissue sections to visualize specific cellular markers in situ. The procedure was as follows: Deparaffinization and rehydration: Slides with FFPE sections were passed through three changes of xylene (5 min each) to remove paraffin. The tissue was then rehydrated through a graded ethanol series (100% ethanol for 3 min, repeated; 95% ethanol for 3 min; 70% ethanol for 3 min) and finally rinsed in distilled water.

Antigen retrieval: Heat-induced epitope retrieval was performed by incubating slides in citrate buffer (pH 6.0) at near-boiling temperature (using a pressure cooker or water bath) for 20–30 min, then allowing them to cool to room temperature.

Blocking: To prevent nonspecific antibody binding, sections were incubated with normal horse serum or normal goat serum (Vector Laboratories) for 30 min at room temperature. The blocking serum was chosen according to the species of the secondary antibody (horse serum when using horse anti-mouse secondary; goat serum when using goat anti-rabbit secondary).

Primary antibody incubation: Sections were incubated for 1 h at room temperature with primary monoclonal antibodies (see Reagents above for details) diluted in antibody diluent. We applied antibodies against targets such as CD3, CD68, TREM2, etc., on serial sections as needed.

Secondary antibody incubation: After a PBS wash, sections were incubated for 30 min with a biotinylated secondary antibody (either horse anti-mouse IgG or goat anti-rabbit IgG from the VECTASTAIN Elite ABC kit, as appropriate for the primary).

Signal amplification: We employed the VECTASTAIN Elite ABC peroxidase system.

Slides were incubated with the pre-formed avidin–biotinylated peroxidase complex (ABC reagent) for 30–60 min at room temperature. The ABC complex binds to the biotin on the secondary antibody, localizing peroxidase at the sites of primary antibody binding.

Chromogen development: Sections were then treated with the substrate solution from the AEC kit (3-amino-9-ethylcarbazole, Vector Laboratories) for approximately 10 min. This reaction deposits a red precipitate at sites of peroxidase activity. The development time was optimized under a microscope to achieve a specific signal with minimal background. The reaction was stopped by rinsing slides in water.

Counterstaining and mounting: Slides were briefly counterstained with hematoxylin to visualize nuclei, then rinsed in running tap water and blued if necessary. Finally, sections were mounted with an aqueous mounting medium (VectaMount AQ, Vector Laboratories) and cover slipped.

Stained sections were examined under a Leica brightfield microscope, and representative images were captured. Whole-slide scans were also acquired using the Motic EasyScan system for high-resolution archival and analysis.

### Tissue immunofluorescence labeling

For immunofluorescent staining of FFPE tissue sections, we followed a protocol similar to the chromogenic IHC through the blocking step, with modifications for fluorescent detection:

Deparaffinization, rehydration, and antigen retrieval: Slides were deparaffinized in xylene (3 × 5 min), rehydrated through graded alcohols to water, and subjected to citrate buffer antigen retrieval (pH 6.0, 20–30 min heat). Slides were cooled and then blocked with normal serum as described above (30 min).

Primary antibody incubation: Sections were incubated for 1 h at room temperature with a mixture of primary antibodies raised in different species or isotypes (to enable multiplexing with distinct secondary antibodies). For example, we combined mouse IgG1, mouse IgG2a, mouse IgG3, and rabbit IgG primary antibodies targeting different antigens of interest. Adjacent sections were incubated with appropriate isotype-matched control antibodies as negative controls.

Secondary antibody incubation: After PBS washes, slides were incubated for 1 h with a panel of fluorescently labeled secondary antibodies chosen to specifically recognize each primary antibody type without cross-reactivity. We used goat anti-mouse IgG subclass-specific antibodies conjugated to Alexa Fluor dyes (e.g., 488, 568, 647) and goat anti-rabbit IgG conjugated to another Alexa Fluor dye. This allowed simultaneous visualization of multiple markers.

Special case – directly labeled antibody: For certain markers, we employed directly conjugated primary antibodies to avoid secondary overlap. For example, in co-localization studies of TREM2 with mycobacterial LAM within lesions, we used an Alexa 488–conjugated anti-TREM2 primary antibody. In such cases, the directly labeled antibody was applied after the unlabeled primaries and their secondaries were completed, to prevent the anti-mouse secondary from binding the labeled TREM2 antibody. We also included an Fc receptor blocking step before adding the direct antibody to minimize any Fc-mediated binding. The Alexa 488–TREM2 antibody was incubated alone for 1 h at room temperature.

Mounting: After final washes (in PBS), sections were mounted using ProLong Gold antifade mounting medium with DAPI (Thermo Fisher), which counterstains nuclei and preserves fluorescence. Coverslips were applied and slides cured overnight before imaging.

High-resolution confocal microscopy was used to image the fluorescently labeled sections. We acquired images on a Leica TCS SP8 MP inverted confocal microscope equipped with 405 nm, 488 nm, 561 nm, and 647 nm lasers. Images were taken under identical settings for samples and controls. Co-localization of fluorescent signals (e.g., TREM2 with LAM) was quantified using ImageJ: we utilized the Colocalization Threshold plugin to calculate the percentage of each signal overlapping in merged channels, confirming genuine co-localization in situ.

### Live *Mtb* and *Mtb* ligands

Virulent *Mtb* H37Rv (ATCC strain) was cultivated for in vitro infection experiments. Bacteria were grown in Middlebrook 7H9 broth (Becton Dickinson) supplemented with 1% glycerol and 10% OADC (oleic acid–albumin–dextrose–catalase; Becton Dickinson) at 37°C in 5% CO₂ (biosafety level 3 conditions). Tween-20 was intentionally omitted from the culture medium to preserve bacterial virulence and prevent washing out of phthiocerol dimycocerosates (PDIM), which are critical lipid virulence factors^57, 85, 86, 87^. This culture methodology has been previously established and used in prior publications from our group^33, 34, 61^. Log-phase cultures were aliquoted in PBS + 10% glycerol and cryopreserved at –80°C. Prior to use, a frozen vial was thawed and the titer verified by plating for colony-forming units (CFU) on Middlebrook 7H11 agar.

We also utilized two engineered fluorescent *Mtb* strains derived from H37Rv6. Strain *Mtb*-BB84 constitutively expresses mCherry, allowing visualization of all bacteria. Strain *Mtb*-LD harbors a dual fluorescent “live–dead” reporter plasmid: it constitutively expresses mCherry and can be induced to express GFP under a tetracycline-inducible promoter system^88^. Both strains were grown in 7H9 medium under the same conditions as wild-type H37Rv, with appropriate antibiotics to maintain plasmids (*Mtb*-LD cultured with hygromycin 50 µg/mL; *Mtb*-BB84 with kanamycin 25 µg/mL). To induce GFP in *Mtb*-LD, we added anhydrotetracycline (ATC, 20 ng/mL) to the culture for 12–24 h prior to use. Successful GFP induction was confirmed by fluorescence microscopy and flow cytometry.

A panel of purified *Mtb* cell envelope ligands was obtained (BEI Resources, NIAID). This included: whole-cell lysate (WCL) of H37Rv, total mycobacterial lipids, phthiocerol dimycocerosate (PDIM), purified mycolic acids, purified lipoarabinomannan (LAM), and purified peptidoglycan. Each ligand was prepared and stored per supplier recommendations (e.g., reconstitution in appropriate solvents). These components were used to stimulate macrophages in vitro (see below).

### Generation of monocyte-derived macrophages and TREM2⁺ macrophages

Primary human monocyte-derived macrophages (MDMs) were generated from peripheral blood mononuclear cells (PBMCs) of healthy adult donors. PBMCs were isolated from whole blood by Ficoll-Paque density gradient centrifugation. CD14^+^ monocytes were then purified using magnetic anti-CD14 microbeads (Miltenyi Biotec) according to the manufacturer’s protocol. The isolated monocytes (typically > 90% purity by CD14 staining) were plated in RPMI 1640 medium with 10% fetal bovine serum (FBS) and human M-CSF (50 ng/mL) to promote differentiation. Cells were cultured at 37°C in 5% CO2, with fresh medium changes every 2–3 days. By day 5–6 of culture, the monocytes had differentiated into adherent MDMs.

Only donors in which ≥80% of macrophages expressed TREM2 following IL-4 stimulation, as determined by flow cytometry, were included for TREM2⁺ macrophage generation and downstream experiments. To generate macrophages with high TREM2 expression (TREM2⁺ macrophages), we further stimulated a subset of MDMs with IL-4. Specifically, on day 5 of differentiation, IL-4 (50 ng/mL) was added to the culture in the continued presence of M-CSF and cells were incubated for an additional 48 h. IL-4 treatment is known to drive an alternative activation state and, in our hands, upregulated surface TREM2 on macrophages. Control MDMs were maintained in M-CSF alone (no IL-4) for the same period. After 2 days, IL-4–treated macrophages were confirmed to have elevated TREM2 surface levels (by flow cytometry), and these cells are referred to as TREM2⁺ macrophages.

### Flow cytometry

Flow cytometry was used to analyze macrophage surface markers under various stimulation conditions. For experiments with live *Mtb* and *Mtb* ligands, we first generated MDMs as described above. On day 5–6 of MDM differentiation, cells were subjected to one of the following treatments for 48 h: (a) IL-4 (100 ng/mL) to induce TREM2, (b) live virulent *Mtb* H37Rv infection at a multiplicity of infection (MOI) of 5 or 10, or (c) stimulation with purified *Mtb* ligands (10 µg/mL each of WCL, total lipids, PDIM, mycolic acids, LAM, or peptidoglycan). After 2 days of stimulation or infection, we harvested the cells for flow cytometric staining. Adherent macrophages were lifted using cold PBS with 1 mM EDTA, collected, and washed in FACS buffer (PBS + 2% FBS). Cells were incubated with fluorochrome-conjugated antibodies against TREM2 (APC anti-TREM2) and CD163 (PerCP-eFluor 710 anti-CD163) on ice for 30 min. Matched isotype control antibodies were included in parallel to assess background staining. After staining, cells were washed and fixed in 1% paraformaldehyde before analysis.

In a parallel set of experiments, we examined the effect of active vitamin D (calcitriol) on TREM2 expression. TREM2+ macrophages (IL-4–treated MDMs) and control MDMs (no IL-4) were treated with 1,25-dihydroxyvitamin D3 (1,25(OH)2D3; 1 × 10−8 M) for 24 h. Following this treatment, cells were harvested and stained for TREM2 as above. This allowed us to determine if calcitriol modulates TREM2 levels on macrophages.

To evaluate macrophage viability following *Mtb* infection, TREM2⁺ macrophages (IL-4–treated MDMs) and control TREM2⁻ MDMs (no IL-4) were infected with GFP-expressing *Mtb* under the conditions described above. After infection, cells were harvested and analyzed by flow cytometry. The gating strategy included exclusion of debris based on forward and side scatter, doublet discrimination (FSC-A vs. FSC-H), and selection of viable cells using a fixable viability dye. Macrophages were then separated into TREM2⁻ and TREM2⁺ populations based on surface TREM2 staining. Infection was determined by detection of GFP signal from live *Mtb*. Data are presented as the percentage of macrophages harboring GFP⁺ *Mtb* within the TREM2⁻ and TREM2⁺ gates. Paired analyses were performed comparing infected cells within each subset.

Stained cells were acquired on a Thermo Fisher Attune NxT flow cytometer operated at the UCLA Flow Cytometry Core Facility. Machine settings (voltages, compensation) were consistent across samples, and at least 10,000 events were collected per condition. Data were analyzed using FlowJo software (v10; Tree Star). Live cell singlets were gated based on forward/side scatter, and fluorescence gates were set using isotype controls. We quantified the percentage of TREM2^+^ and CD163^+^ cells, as well as median fluorescence intensity, under each condition to assess macrophage phenotypic changes in response to stimuli.

### TREM2 activation assay (TREM2 reporter)

We utilized a cell-based reporter system to test whether *Mtb* and its components can directly trigger TREM2–DAP12 signaling. The reporter cells, known as 2B4-DAP12-GFP cells, are an engineered murine T cell hybridoma (2B4 line) expressing human TREM2 along with the DAP12 signaling adaptor^22, 89^. In these cells, engagement of TREM2 leads to DAP12 phosphorylation and downstream activation of an NFAT-responsive GFP reporter, such that GFP expression serves as a proxy for TREM2 activation.

For the assay, 2B4-DAP12-GFP/TREM2 cells were plated in 96-well plates coated with various stimuli and then incubated at 37°C for 24 h. The stimulation conditions included: (i) plate-bound anti-TREM2 monoclonal antibody (positive control for maximal TREM2 cross-linking), (ii) soluble mycobacterial ligands (e.g., LAM, mycolic acids, etc., added into the culture medium), and (iii) plate-bound mycobacterial lipid extracts (to mimic presentation of cell wall lipids in a solid-phase context). For plate-coated conditions, high-binding plates were pre-coated overnight at 4°C with either 5 µg/mL anti-TREM2 antibody or 10 µg/mL of purified lipid, then washed before adding cells.

After 24 h of stimulation, we gently resuspended the cells and measured GFP expression by flow cytometry. The cells were first evaluated under a fluorescence microscope to qualitatively confirm GFP induction. Then they were analyzed on an Attune NxT flow cytometer (Thermo Fisher) to quantify the percentage of GFP-positive cells or the mean fluorescence intensity of GFP. Unstimulated reporter cells and cells with an isotype antibody coating served as negative controls to establish baseline GFP levels. An increase in GFP signal in cells exposed to *Mtb* components (relative to controls) indicated activation of TREM2–DAP12 signaling by those components.

### Detection of lipid droplets in macrophages

TREM2^+^ macrophages and control MDMs were cultured on chambered coverglass slides (2 × 10^5 cells per well) and subjected to various treatments for 24 h to induce lipid droplet accumulation: (1) IL-4 (100 ng/mL) alone, (2) live mCherry-expressing *Mtb* (strain *Mtb*-BB84, MOI 5), and (3) a cocktail of *Mtb* cell envelope ligands (10 µg/mL each, as detailed above).

#### A. Oil Red O Staining

Oil Red O (Sigma-Aldrich, Cat. No. O0625) was used to stain neutral lipids (lipid droplets) in human monocyte-derived macrophages (MDMs). After stimulation, cells were fixed in 10% neutral-buffered formalin for 30 minutes, rinsed in distilled water, and equilibrated in 60% isopropanol. Lipid droplets were stained with freshly prepared ORO working solution (3:2 stock: water) for 15 minutes, followed by hematoxylin counterstaining for 2 minutes. Slides were mounted in aqueous medium and scanned using a Motic EasyScan digital slide scanner. Lipid burden was quantified in ImageJ by measuring the ORO-positive area per cell and expressed as fold change relative to untreated controls.

#### B. SMCy5.5 fluorescent lipid dye

We examined lipid droplet formation in macrophages in response to *Mtb* using the fluorescent dye SMCy5.5 (TOCRIS Bioscience), which specifically stains neutral lipid droplets. SMCy5.5 is a lipophilic cyanine dye with far-red emission (∼680 nm) that brightly labels lipid storage organelles while remaining compatible with cell fixation. A 5 mM stock solution of SMCy5.5 in DMSO was prepared and stored light-protected at –20°C.

After the 24 h stimulation, we applied SMCy5.5 to stain intracellular lipids. Cells were gently washed with warm PBS to remove extracellular bacteria and debris, then incubated with 1 µM SMCy5.5 in serum-free, phenol red–free RPMI 1640 for 30 min at 37°C. (In preliminary tests, a 30–60 min staining period was sufficient for clear droplet labeling.) After staining, cells were washed with PBS and fixed in 4% paraformaldehyde for 30 min at room temperature. The chambers were removed, and slides were mounted with ProLong Diamond antifade mountant containing DAPI (to counterstain nuclei).

Fluorescent imaging was performed using a Leica TCS SP8 confocal microscope. We used a 633 nm laser line to excite SMCy5.5 and collected emission in the far-red channel; DAPI nuclear staining was imaged in the blue channel. Z-stack images were acquired to ensure capture of droplets throughout the cell volume. Image analysis confirmed the presence of bright cytoplasmic lipid droplets in *Mtb*-infected macrophages, particularly in the IL-4–polarized (TREM2^+)^ macrophages. We qualitatively noted increased lipid droplet formation in infected TREM2^+^ macrophages compared to controls, consistent with a foamy macrophage phenotype.

### *Mtb*–GFP–mCherry infection and viability assay

We used a dual-fluorescent *Mtb* reporter strain to simultaneously assess bacterial uptake (infection rate) and viability inside macrophages. The reporter strain (LD-*Mtb*, derived from H37Rv) constitutively expresses a red fluorescent protein (mCherry) and conditionally expresses green fluorescent protein (GFP) under a tetracycline-inducible promoter6. In this system, all bacteria fluoresce red, but only metabolically active (viable) bacteria can induce GFP expression upon addition of the inducer (ATC). Thus, within infected cells, red-only bacteria are presumed dormant or non-viable, whereas red+green bacteria are viable.

TREM2^+^ macrophages and control MDMs were seeded in 8-well chamber slides (2 × 10^5^ cells per well) and allowed to adhere. For baseline infection experiments, macrophages were infected with LD-*Mtb* at MOI 5. The bacteria (pre-induced with ATC so that any viable bacteria would be GFP+) were added to the cells and allowed to infect for 24 h at 37°C. After infection, cells were washed to remove extracellular bacilli and then fixed with 4% PFA. Slides were mounted with ProLong Diamond containing DAPI.

For the vitamin D3 modulation experiment, TREM2^+^ and control macrophages were pre-treated with 1,25(OH)2D3 (1 × 10−7 M, approximately 100 nM) for 24 h. After this pre-treatment, the macrophages were infected with the LD-*Mtb* reporter (MOI 5) for an additional 24 h, then processed similarly for imaging. This experiment was designed to test whether calcitriol activation of macrophages could enhance their antimicrobial activity against *Mtb*.

Confocal microscopy was used to image infected cells. mCherry fluorescence (red) marked all *Mtb* present, and GFP fluorescence (green) marked viable, transcriptionally active *Mtb*. We captured multiple fields for each condition and donor. Infectivity was quantified by measuring the total bacterial burden per cell: we used ImageJ to quantify the integrated density of mCherry signal in at least 30 macrophages per condition (from multiple images) and calculated the average mCherry intensity per cell. This reflects the average number of bacilli per macrophage.

Bacterial viability was quantified by assessing the co-localization of GFP and mCherry signals within each cell. For each analyzed macrophage (≥ 30 cells per condition), we determined the fraction of the mCherry+ bacterial area that was also GFP+ (indicating live bacteria). We utilized the ImageJ Colocalization Threshold plugin for this analysis, which computes the degree of pixel-wise overlap between the two fluorescent channels after automatically thresholding each. We reported the GFP–mCherry co-localization as a percentage of total *Mtb* (mCherry) signal per cell. Macrophages containing GFP+ *Mtb* were interpreted as harboring live bacteria. By comparing this percentage between TREM2+ vs control macrophages and ± vitamin D treatment, we evaluated the impact of TREM2 polarization and vitamin D on the control of intracellular *Mtb* viability.

### LC3 Immunofluorescence Labeling

LC3 immunofluorescence was performed to quantify autophagy activation in MDMs and TREM2⁺ macrophages during *Mtb* infection. TREM2⁺ macrophages and control MDMs were seeded into 8-well chamber slides (2 × 10⁵ cells per well) and allowed to adhere overnight. Cells were infected with LD-*Mtb* at an MOI of 5; bacteria were pre-induced with anhydrotetracycline (ATC) to ensure GFP expression in viable bacilli. Following 24 hours of infection at 37 °C, cells were washed thoroughly with warm PBS to remove extracellular bacteria and fixed in 4% paraformaldehyde for 20 minutes at room temperature. Fixed cells were permeabilized using Cytofix/Cytoperm™ solution (BD Biosciences) for 30 minutes, blocked with serum for 20 minutes, and incubated with anti-LC3 primary antibody for 1 hour at room temperature. After washing, cells were incubated with Alexa Fluor 647-conjugated secondary antibody for 60 minutes, washed again, and mounted using ProLong Gold antifade reagent with DAPI. Slides were cured overnight before imaging.

Immunofluorescence images were acquired using a Leica TCS SP8 MP inverted single-photon confocal laser-scanning microscope (Leica Microsystems, Heidelberg, Germany) at the Advanced Microscopy/Spectroscopy and Macro-Scale Imaging Laboratory of the California NanoSystems Institute, University of California, Los Angeles. Identical acquisition settings were used across donors and conditions. LC3 puncta were quantified using ImageJ and expressed as the number of LC3⁺ puncta per cell.

### Quantification of *Mtb* growth by CFU assay

To directly measure the ability of macrophages to restrict or kill intracellular *Mtb*, we performed colony-forming unit (CFU) assays at multiple time points. TREM2^+^ macrophages and control MDMs were first treated with 1,25(OH)_2_D_3_ (10^−7^ µM, 24 h) or left untreated. Cells were then infected with virulent H37Rv *Mtb* at MOI 5. We evaluated bacterial CFUs at two time points: an initial time point shortly after infection (Day 0, defined as 2 h post-inoculation, after washing off extracellular bacteria) and after 4 days of incubation (Day 4).

At each time point, infected macrophages were lysed to release intracellular *Mtb*. We lysed cells by adding 0.3% saponin in PBS for 10 min at 37°C, which effectively solubilizes the host cell membrane without killing *Mtb*. Lysates were pipetted up and down to disrupt cell clumps and then transferred to sterile tubes. To further break up any bacterial aggregates, we sonicated the lysates for 5 min in a bath sonicator set to 37°C. Serial dilutions of the lysates were prepared in PBS. These dilutions were plated on Middlebrook 7H11 agar plates supplemented with OADC and glycerol. Plates were incubated at 37°C and monitored for Mtb colony growth.

After 3 weeks, colonies were counted on plates from appropriate dilution levels. CFU counts (per well or per initial cell number) were calculated for Day 0 and Day 4 samples. The ratio of Day 4 to Day 0 CFUs provided a measure of *Mtb* growth (or killing) inside the macrophages. We compared CFU outcomes between TREM2^+^ vs control macrophages and ± vitamin D treatment. Consistently lower CFUs in TREM2^+^ or 1,25(OH)_2_D_3_ –treated groups would indicate enhanced bacterial killing or growth restriction. These CFU assays complemented the fluorescence viability imaging, offering a quantitative microbiological confirmation of intracellular bacterial survival.

### Statistics for in vitro assays

All quantitative in vitro experiments were performed on biological replicates (macrophages derived from multiple donors), and results are reported as mean ± standard error of the mean (s.e.m.). Statistical analysis was carried out using GraphPad Prism 10 software. For comparisons involving more than two matched groups (e.g., different treatments on the same donor’s cells), we applied repeated-measures one-way analysis of variance (ANOVA). When the assumption of sphericity was violated, the Greenhouse–Geisser correction was used. Post hoc pairwise comparisons were conducted with Holm–Šídák adjusted P values, unless stated otherwise. For experiments with two independent variables (e.g., genotype × treatment), we used two-way ANOVA to test for main effects and interactions, followed by Tukey’s or Sidak’s multiple comparisons tests as appropriate. For direct two-group comparisons, we used two-tailed Student’s t-tests. In all analyses, a P value < 0.05 was considered statistically significant.

### Correlation analysis

To assess the relationship between lipid metabolism, autophagy, and intracellular M*. tuberculosis* viability, we performed Pearson correlation analysis using donor-matched quantitative measurements obtained across four experimental conditions (MDM–Mtb, MDM–*Mtb*+1,25D, TREM2⁺ MØ–*Mtb*, TREM2⁺ MØ–Mtb+1,25D). For each donor, Mtb viability (GFP signal per cell), lipid droplet content (SMCy5.5 fluorescence), and LC3 puncta were quantified, generating 16 paired observations (4 donors × 4 conditions). Pearson correlation coefficients (r) and associated two-tailed P values were calculated using Prism, with P < 0.05 considered statistically significant. To evaluate robustness, nonparametric permutation testing (10,000 permutations) was additionally performed for each pairwise comparison, and 95% confidence intervals for r were computed using bias-corrected bootstrapping. No data transformation or imputation was applied; all correlations were performed on raw, donor-matched values.

## Data and Code Availability

GeoMx transcriptomic data have been deposited in the Gene Expression Omnibus GEO) under accession number GSE301876. The FFPE scRNA-seq data generated in this study have been deposited in the GEO under accession number GSE301610, including sample series GSM9086459. All data supporting the findings of this study are also available at Zenodo (DOI: 10.5281/zenodo.15750734). Custom scripts and analysis pipelines are available on GitHub at https://github.com/modlab246/TREM2_PTB.

## Acknowledgments

We thank Dr. Robert Hunter for providing insight and advice on the lung pathology of tuberculosis (TB) as well as the pathogenesis of TB infection. We are grateful to Christopher C. Tai K. for assistance with single-cell RNA sequencing (scRNA-seq) from FFPE pulmonary TB samples, and to Annaliza Legaspi for help with blood donor selection and processing. We acknowledge the University of California–Los Angeles California NanoSystems Institute Advanced Light Microscopy Core Facility for assistance with confocal studies (RRID: SCR_022789). We thank Barbara Dillon from the UCLA High Containment Facilities for support with BSL-3–required assays. We also thank the UCLA Technology Center for Genomics & Bioinformatics (TCGB) for assistance with GeoMX and scRNA-seq, the Translational Pathology Core Laboratory (TPCL) for sectioning FFPE TB samples for CosMx and scRNA-seq, and the UCLA Jonsson Comprehensive Cancer Center (JCCC) Flow Cytometry Core Facility for support with flow cytometry assays.

## Authors contribution

R.L.M. supervised and conceptualized the study. R.M.B.T. led the experimental work, including functional assays and CosMx, contributed to conceptualization and writing. C.B. contributed to sample collection, conceptualization and writing. J.P. analyzed GeoMX data and contributed to bioinformatics. C.W. and Y.G. analyzed FFPE RNA-seq data. J.W. contributed to CosMx, scRNA-seq, and functional analysis. B.J.A.S. performed *Mtb* infections and CFU assays. P.R.A. conducted RNAscope analysis. L.M., K.L., and F.M. performed bioinformatics and gene signature analyses. P.D. supported CosMx assays. L.F. contributed to mycobacterial ligand assays. M.B. performed Oil red experiments. K.M. and A.P. performed immunohistochemistry. S.M.F., M.R.B., E.R., and S.R. provided pathology support. A.J.C.S. supported RNAscope assays. E.K., M.C., S.B., and D.L.B. supported gene signature analyses. P.S.B., B.D.B., M.P., and J.T.B. contributed to data interpretation and manuscript revision. B.R.B. contributed to conceptualization, data interpretation and writing. The manuscript was written by R.M.B.T., B.R.B. and R.L.M.

