## supplemental figures for "TREM2^+^ macrophages accumulate in alveoli of human pulmonary tuberculosis providing a permissive niche for bacterial growth"

#### Supplemental Figure 1

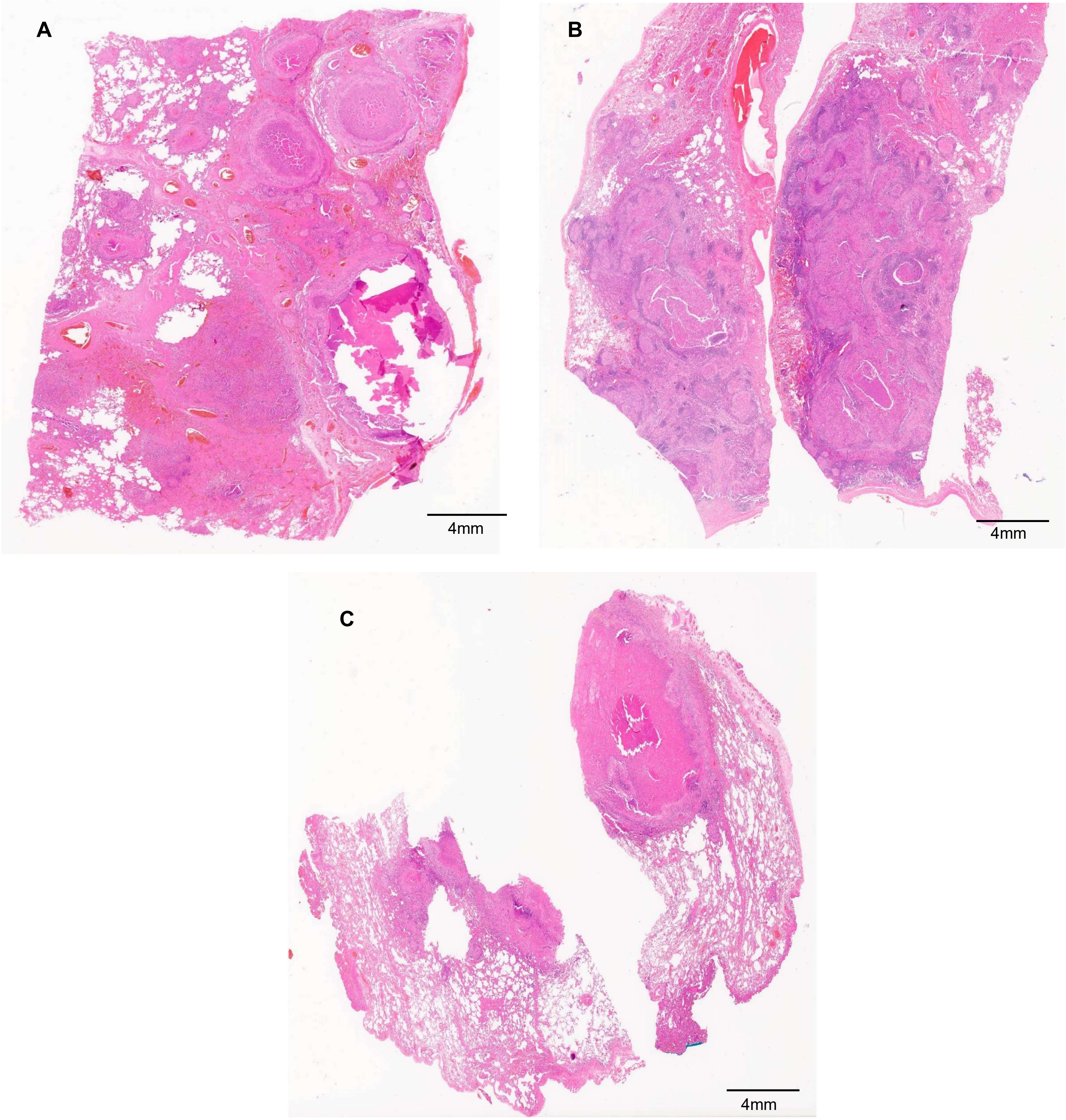


**Supplemental Figure 1**: Histology of pulmonary TB (PTB) specimens used for GeoMx spatial profiling. (A) PTB 22.1; (B) PTB 22.2 and (C) PTB 22.3. H&E.

#### Supplemental Figure 2

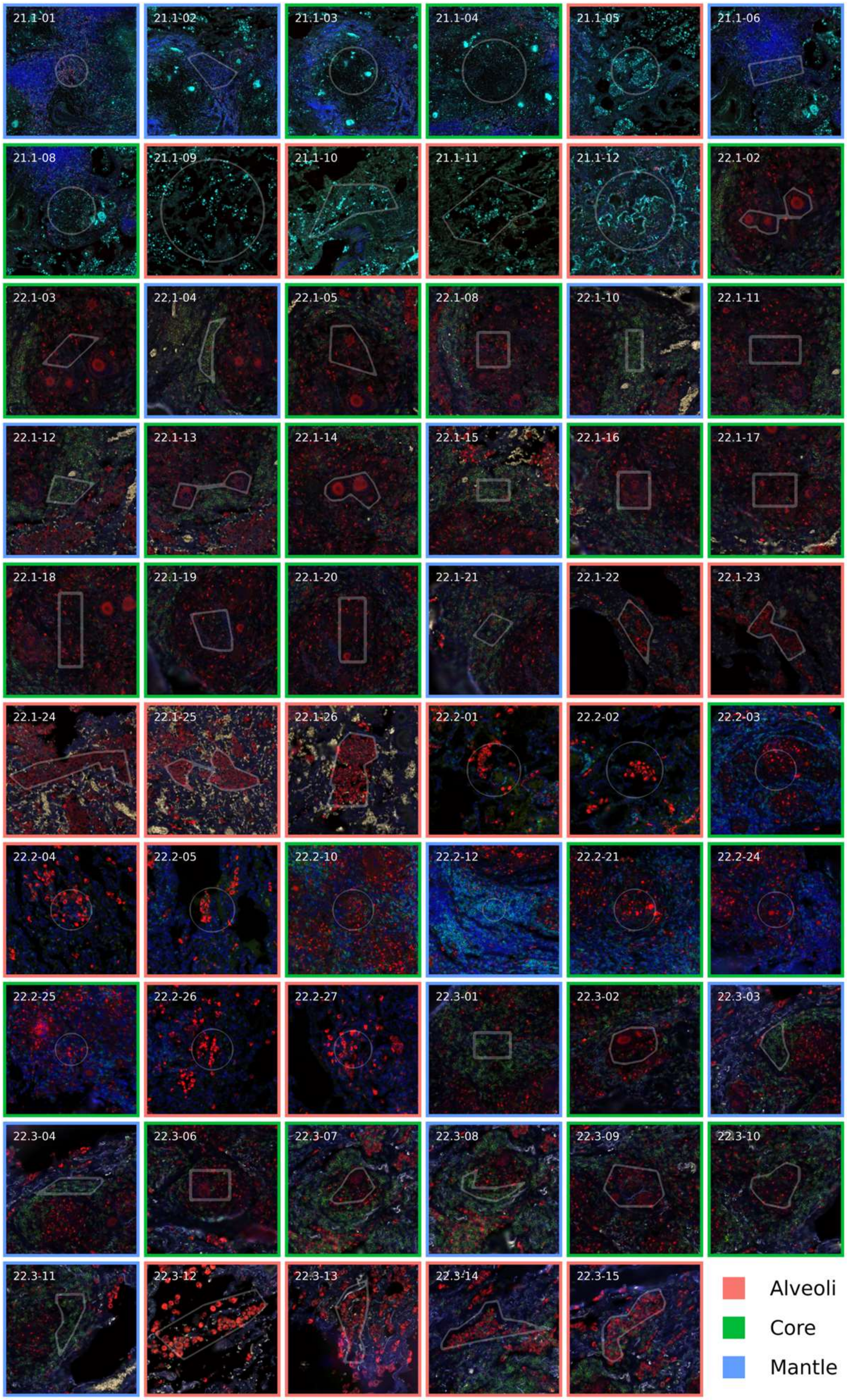


**Supplemental Figure 2. Spatial profiling of pulmonary tuberculosis (PTB) lesions using GeoMx: selection of regions of interest (ROIs).** Representative tissue sections from four pulmonary TB (PTB) cases (PTB 21.1a, PTB 22.1, PTB 22.2, and PTB 22.3) were analyzed using the GeoMx Digital Spatial Profiler. Regions of interest (ROIs) were strategically selected across granulomatous and adjacent lung tissue areas to capture spatial transcriptional heterogeneity associated with TB pathology.

### Supplemental Figure 3

A

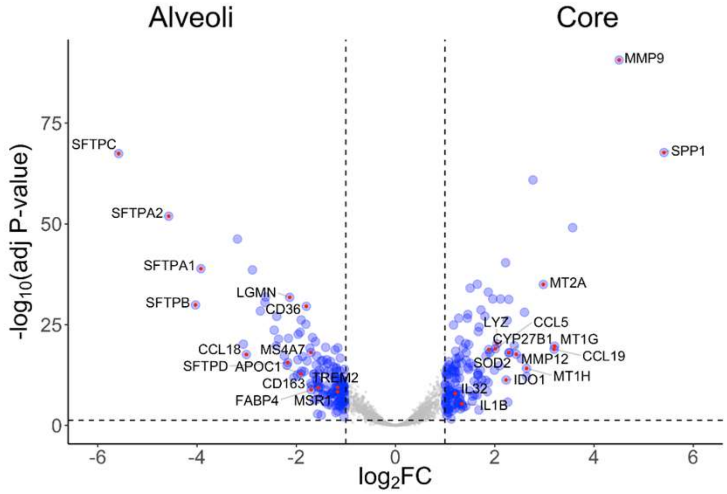


B

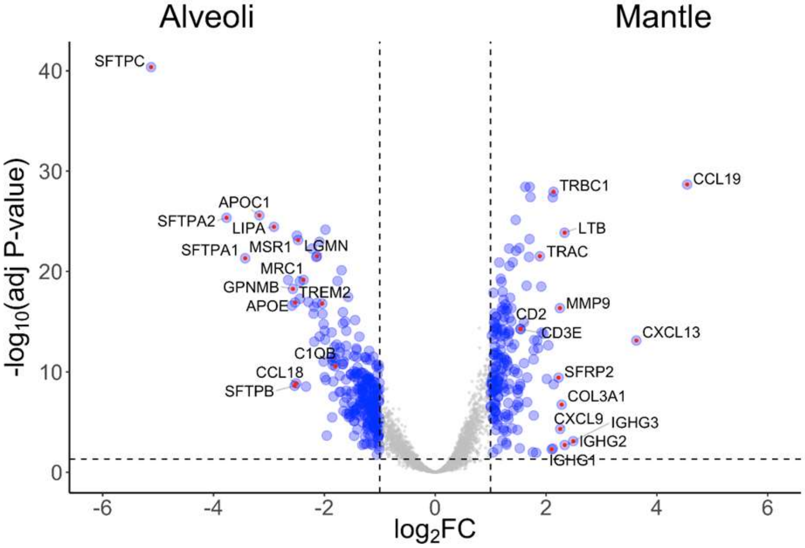


C

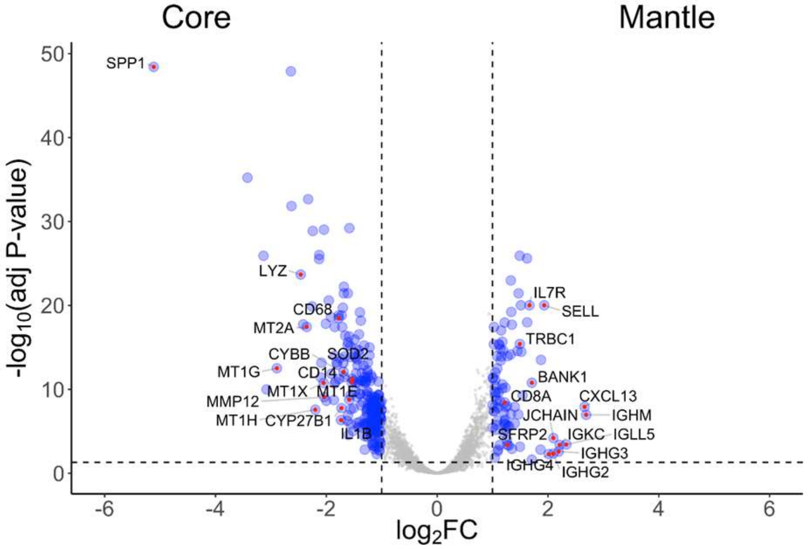


**Supplemental Figure 3.** GeoMx analysis of pulmonary TB specimens. Volcano plots showing log2FC and –log10(padj) for expression of 202 genes in alveoli vs. core (A), alveoli vs. mantle (B) and core vs. mantle (C).

### Supplemental Figure 4

A

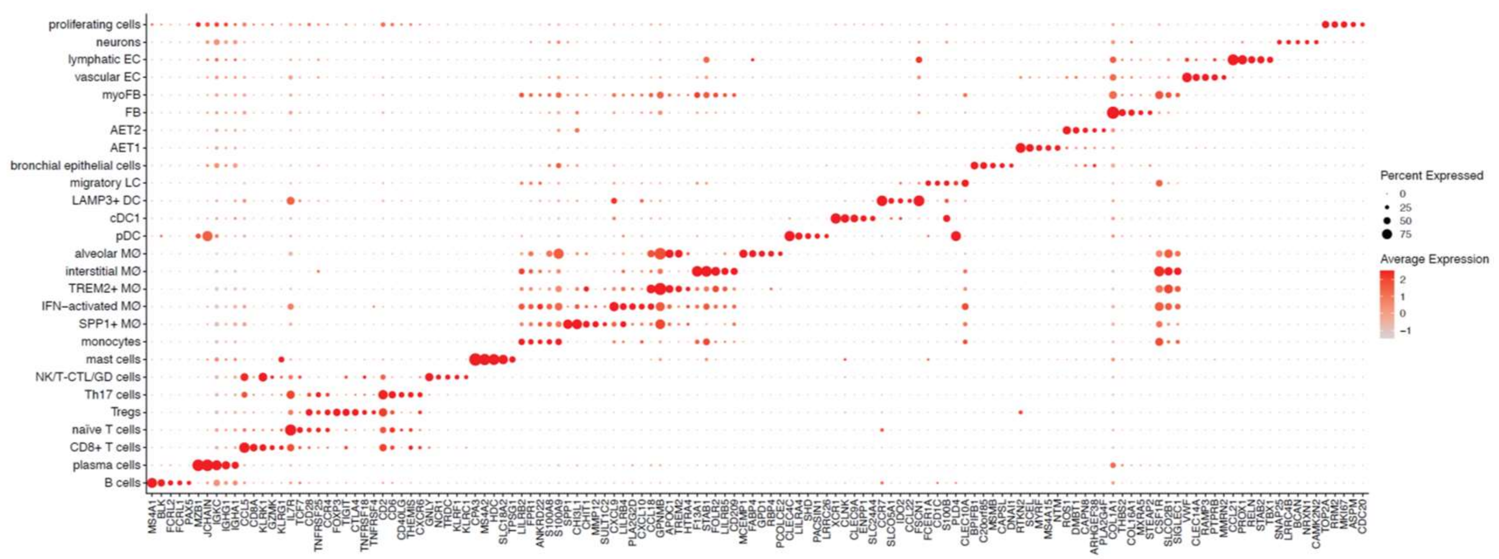


B

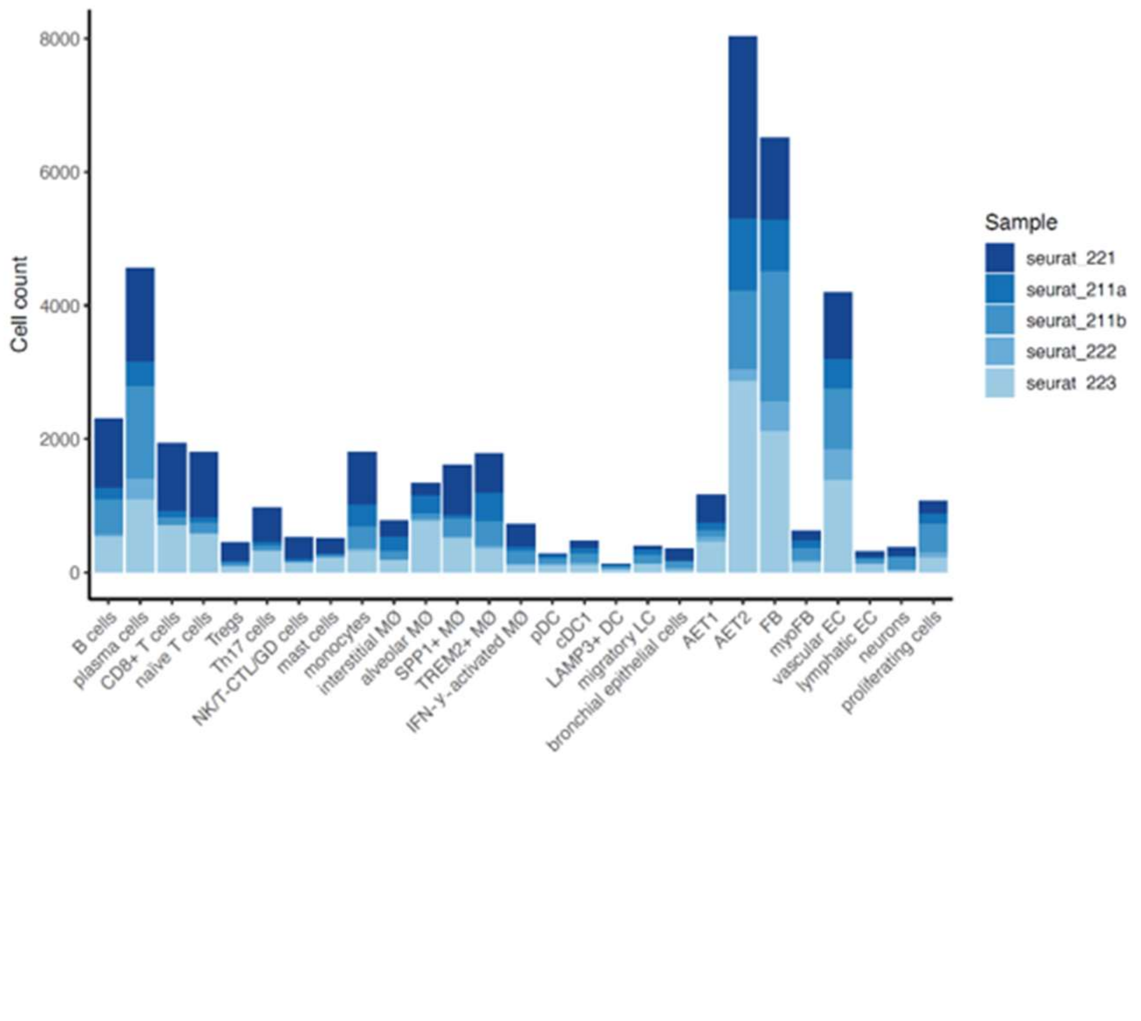


**Supplemental Figure 4.** Cell cluster definitions for PTB scRNA-seq data. The scRNA-seq data from PTB samples were clustered into major cell types, and then the myeloid and T cell clusters were further subclustered. (A) Top five signature genes in each cluster. (B) Number of cells in each cluster by PTB sample number.

#### Supplemental Figure 5

A B

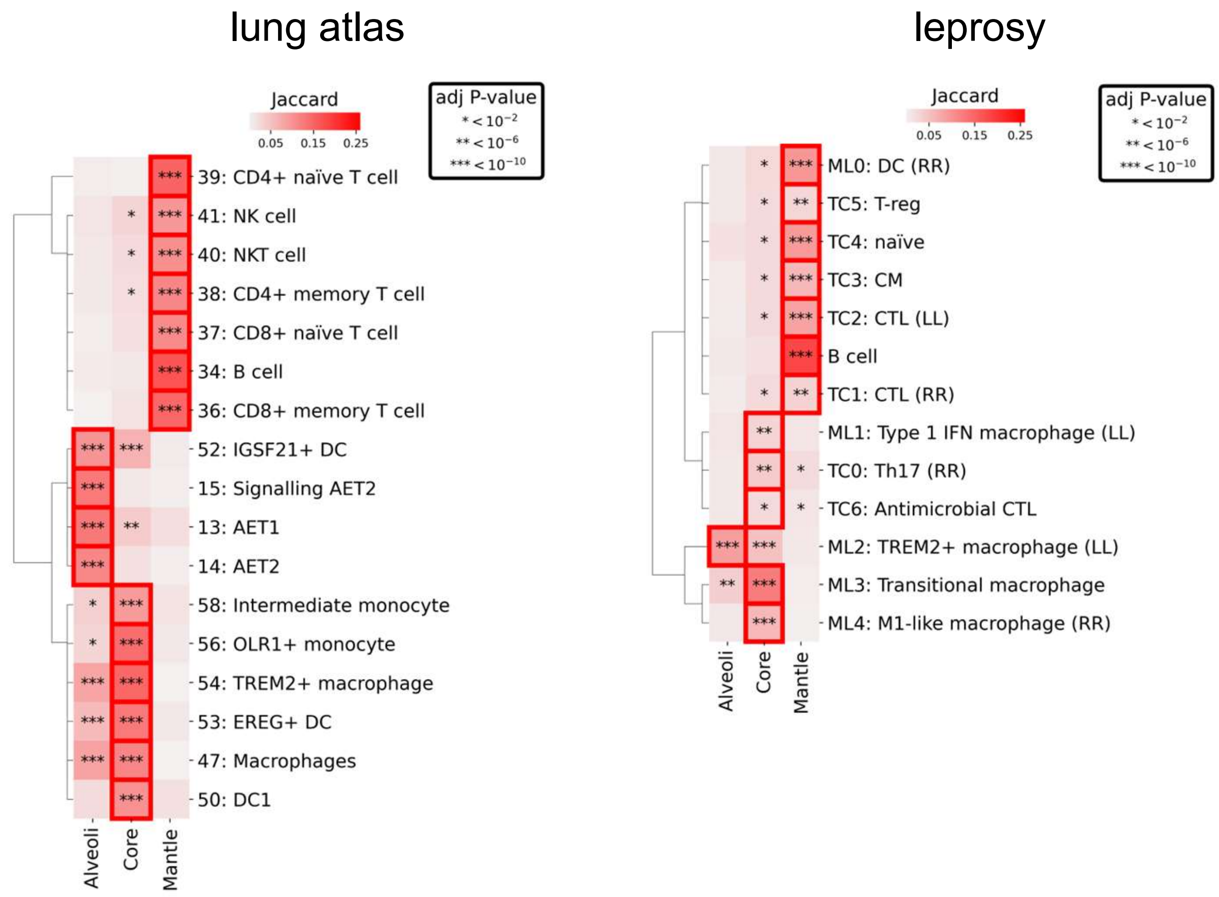


**Supplemental Figure 5.** Integration of GeoMx pulmonary TB (PTB) spatial profiling data with scRNA-seq atlases. (A) Jaccard overlap of GeoMx PTB spatial profiling data with the human lung atlas scRNA-seq data. (B) Jaccard overlap of GeoMx PTB spatial profiling data with the leprosy scRNA-seq data. Hypergeometric distribution analysis was used for statistical analysis.

### Supplemental Figure 6

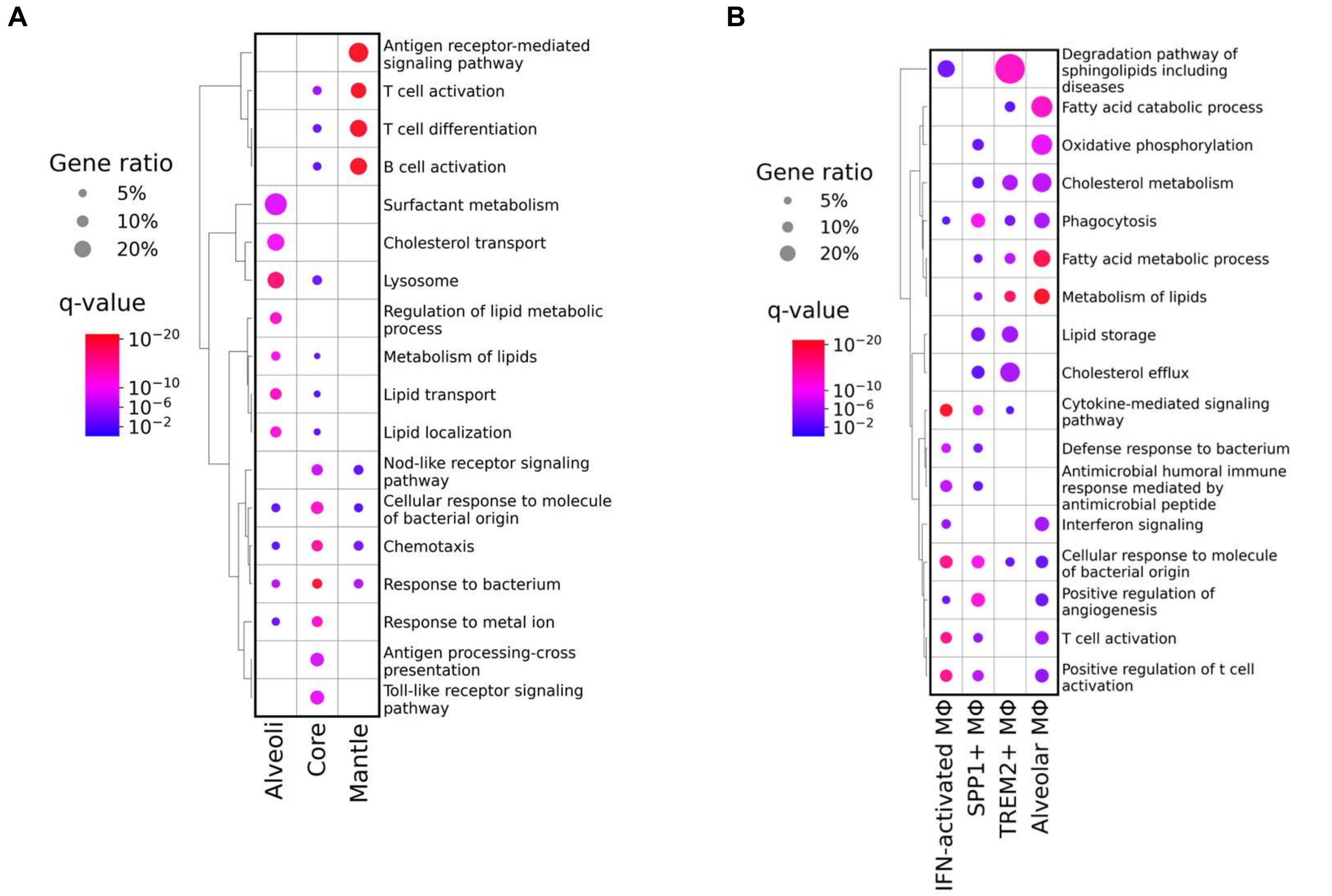


**Supplemental Figure 6**. Functional pathway analysis of the GeoMx spatial profiling data and scRNA-seq data from human pulmonary TB (PTB). (A) Metascape analysis of GeoMx spatial profiling differential expressed genes (DGEs) from PTB to identify functional pathways. (B) Metascape analysis of scRNA-seq signatures from PTB to identify functional pathways.

### Supplemental Figure 7

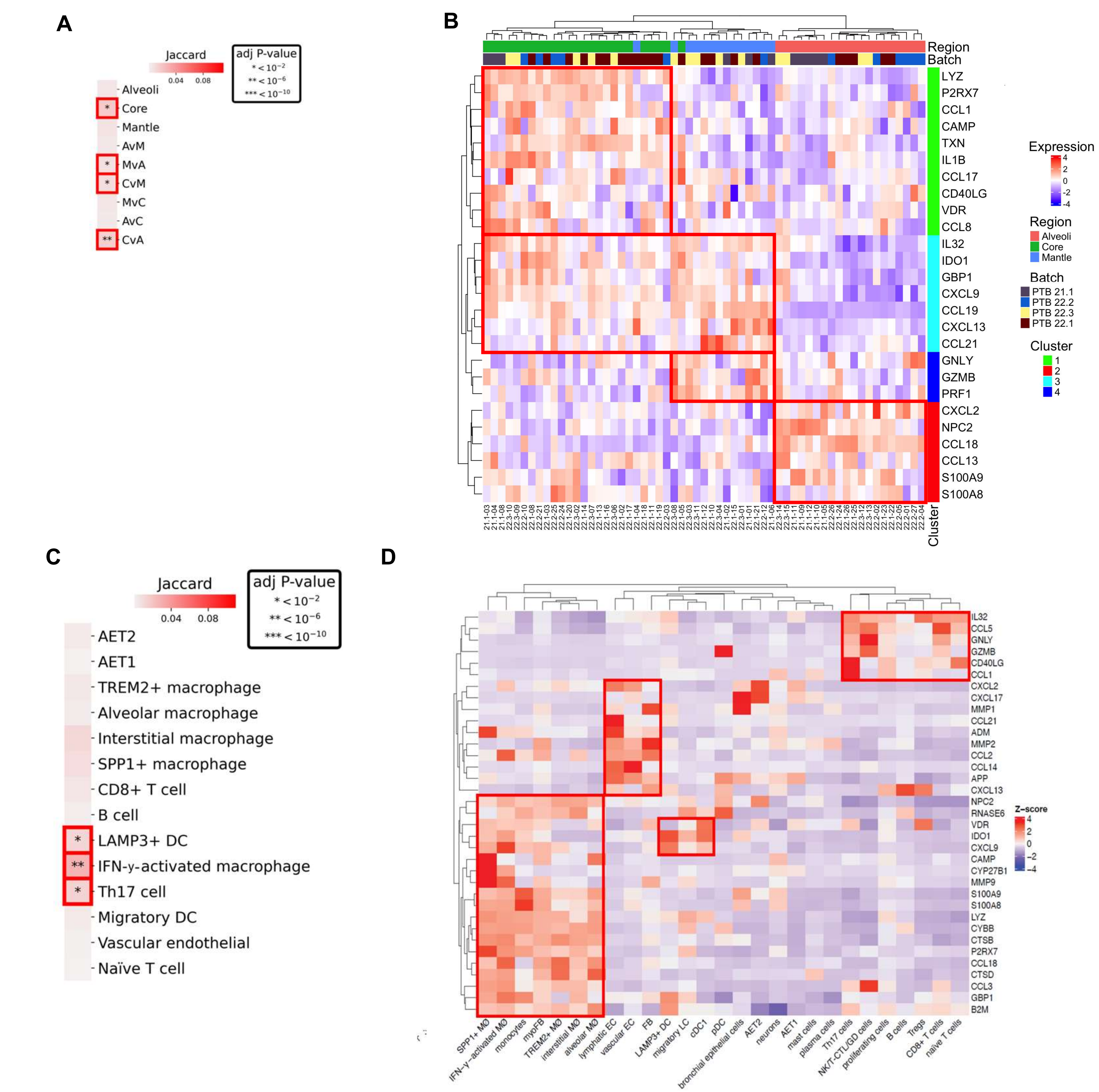


**Supplemental Figure 7.** Antimicrobial genes signature in PTB samples. (A) Antimicrobial genes expressed in alveoli (PTB A), core (PTB C) and mantle (PTB M) from PTB GeoMx data by Jaccard index and hypergeometic distribution analysis. (B) Heatmap of antimicrobial gene expression in alveoli, core and mantle form PTB GeoMx data. (C) Antimicrobial genes expressed by the PTB cell types from scRNA-seq data by Jaccard index and hypergeometric distribution analysis. (D) Heatmap of antimicrobial genes expressed by the PTB cell types from scRNA-seq data.

### Supplemental Figure 8

A

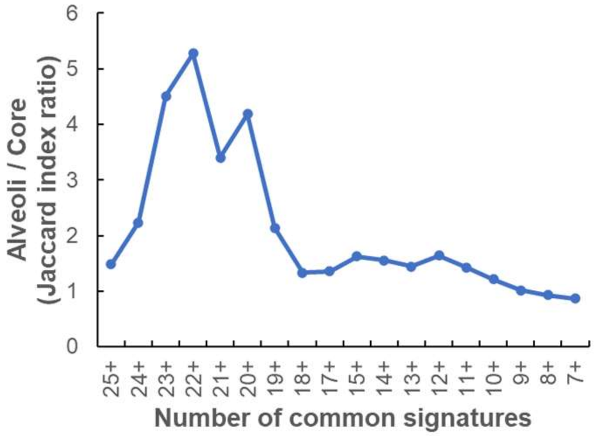


B

| Gene | Frequency |
| --- | --- |
| TREM2 | 31 |
| APOE | 29 |
| CTSB | 27 |
| MS4A6E | 25 |
| C1QC | 24 |
| C1QB | 24 |
| GRN | 23 |
| LGMN | 23 |
| CTSD | 23 |
| C1QA | 23 |
| FTL | 22 |
| NPC2 | 22 |

C

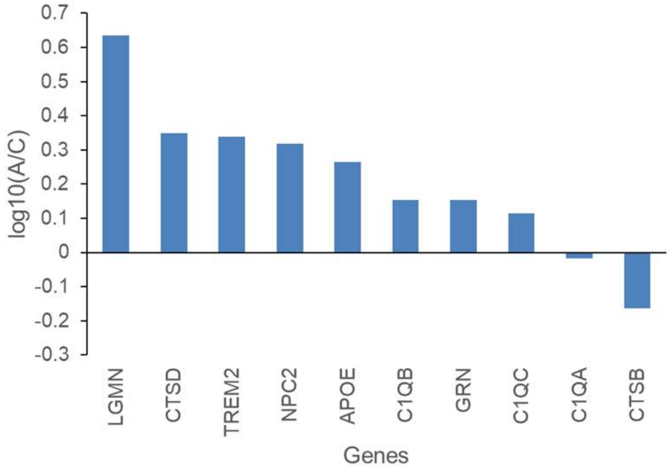


D

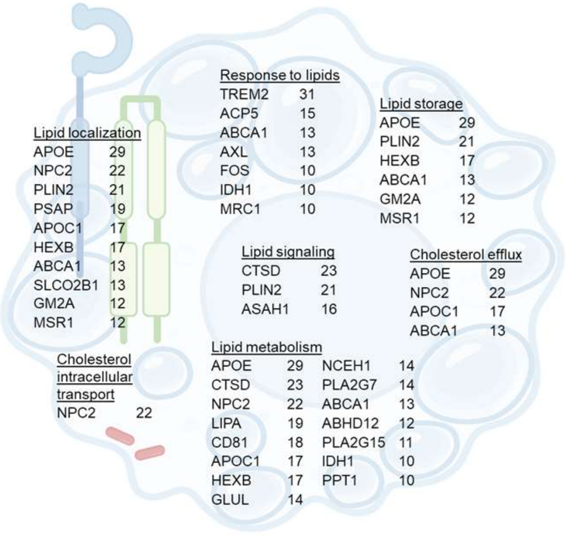


**Supplemental Figure 8**. Expression of alveoli DGEs in TREM2⁺ macrophages. (A) The frequency of individual genes within 34 TREM2⁺ macrophage signatures from 21 publications was calculated and plotted vs. the Jaccard index ratios of these genes in the alveoli/core GeoMX spatial profiling pulmonary TB data. (B) The twelve genes found in 22 or more published TREM2⁺ macrophage signatures are shown with the frequency of that gene in the signatures. (C) The gene expression ratio (log10, alveoli/core) for genes found in 22 or more TREM2 macrophage signatures. MS4A6E and FTL were not detected in the pulmonary TB GeoMx spatial profiling data. (D) TREM2⁺ macrophages show enriched expression of genes involved in lipid metabolism, including lipid localization, signaling, response, and cholesterol transport/efflux.

#### Supplemental Figure 9

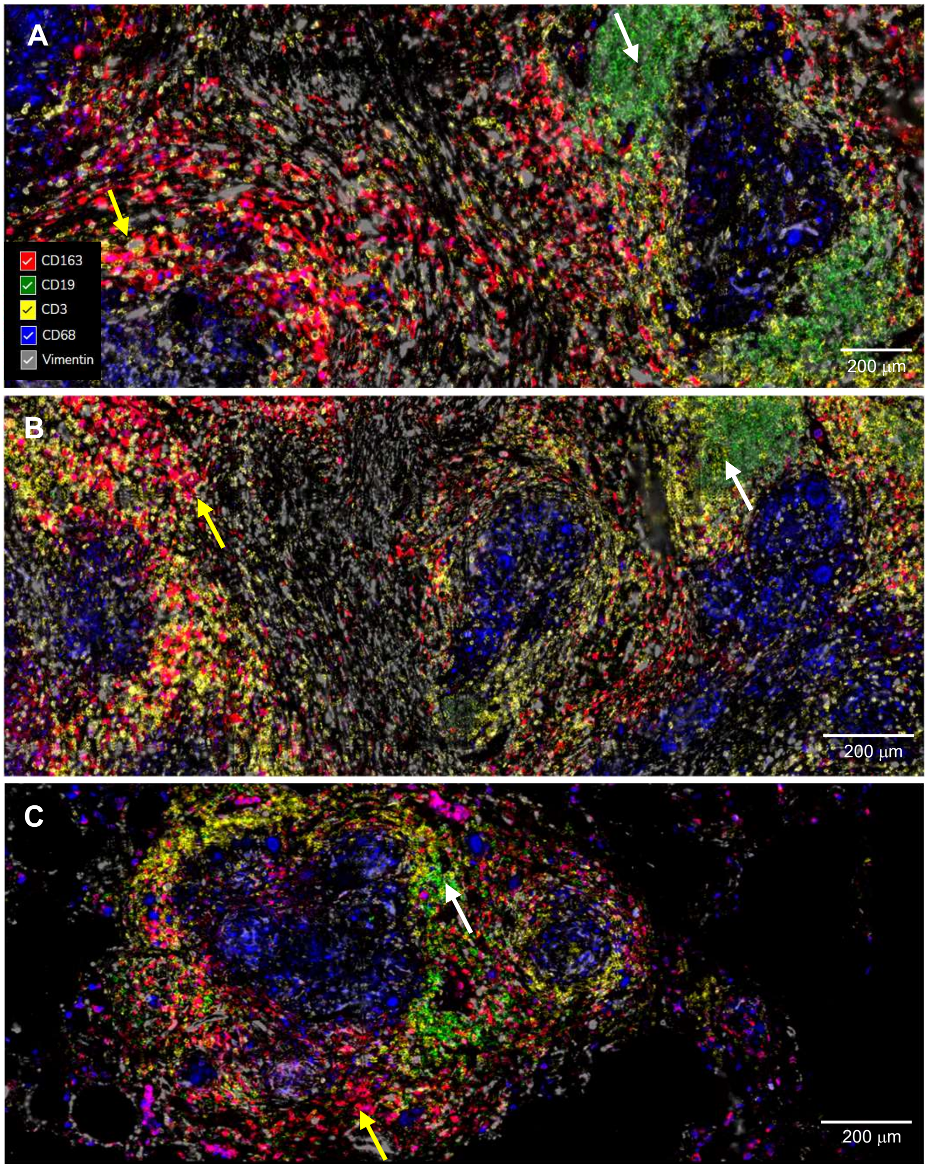


**Supplemental Figure 9**: Immunofluorescence images obtained using the CosMx spatial profiling with the human protein panel. Different types of granulomas in PTB 22.2 (A-B) and PTB22.1 (C) specimens. The granulomas are surrounded by pneumonitis (yellow arrows) and aggregations of B cells (white arrows). Markers: CD163 (red), CD19 (green), CD3 (yellow), CD68 (blue), and vimentin (gray). Scale bars: 200 µm

#### Supplemental Figure 10

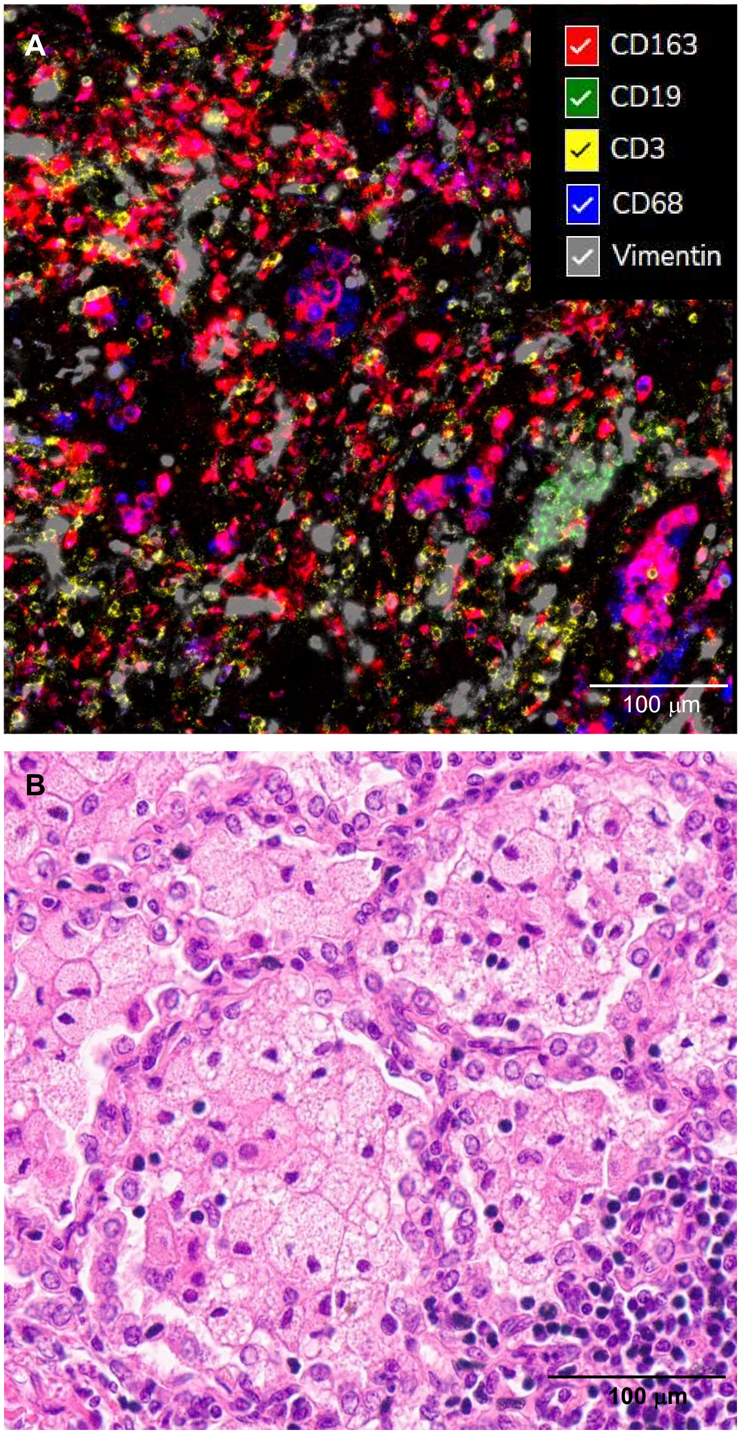


**Supplemental Figure 10. Foamy macrophages in alveoli of pulmonary tuberculosis (PTB).** (A) Immunofluorescence images obtained using the CosMx spatial profiling with the human protein panel, showing CD163+ foamy macrophages (red) surrounded by alveolar epithelium identified by vimentin (gray). Additional markers include CD19 (green), CD3 (yellow), and CD68 (blue). (B) Hematoxylin and eosin (H&E) staining of the same region in the PTB 22.2 specimen. Scale bars: 100 µm.

### Supplemental Figure 11

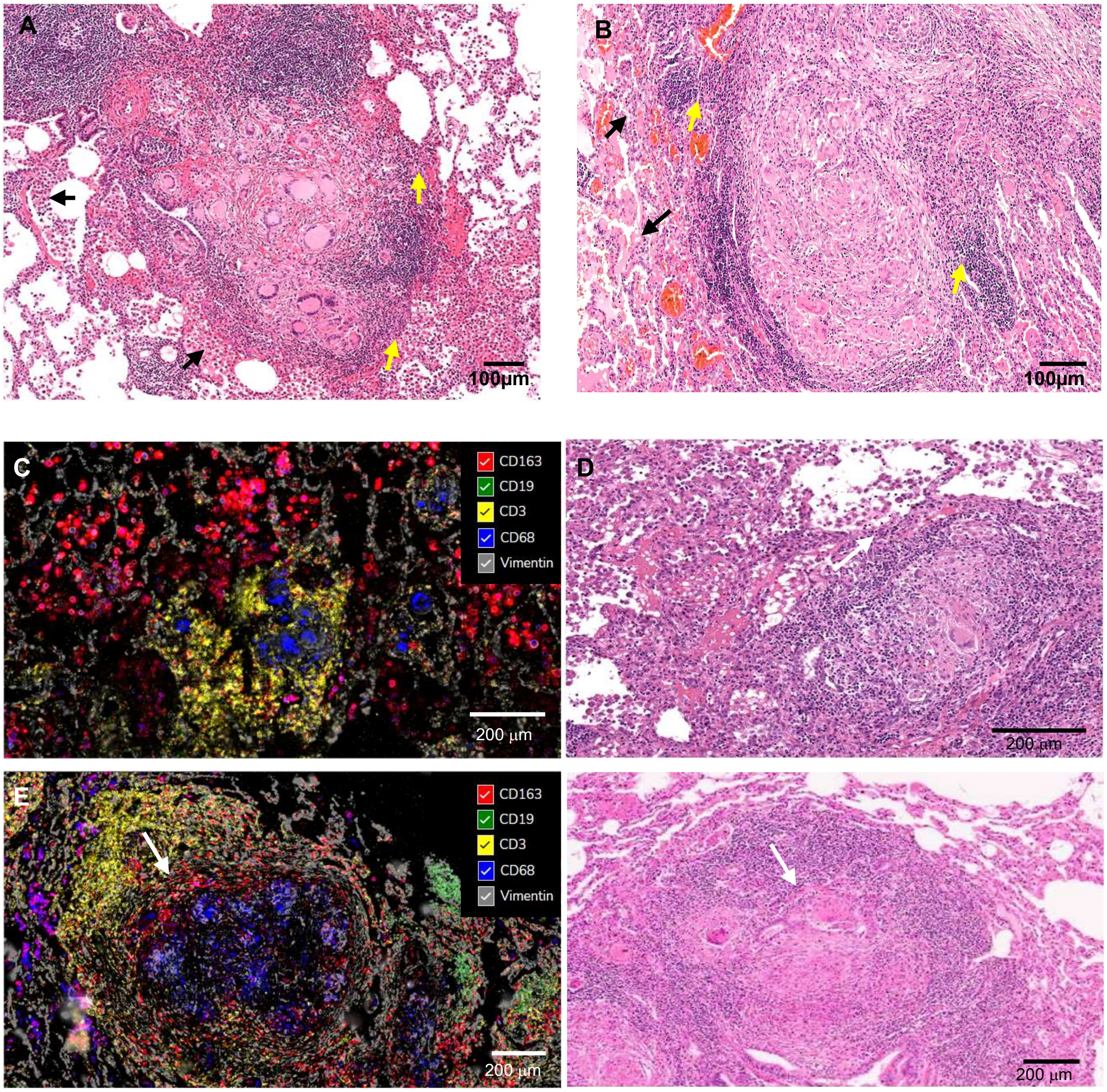


**Supplemental Figure 11**: Histological analysis of pulmonary tuberculosis (PTB) granulomas arising within alveolar spaces. (A, B) Foamy macrophages fill the alveoli (black arrows), while alveolar macrophages are observed in direct contact with the granuloma mantle (yellow arrows) in samples PTB 21.1 (A) and PTB 22.2 (B). (C, D) Organized granulomas adjacent to alveoli exhibit central Langhans giant cells surrounded by a mantle zone rich in T and B lymphocytes. Dense aggregates of T or B cells are noted (PTB 21.1). (E, F) Alveolar macrophages (CD163+, white arrows) are positioned at the interface between the granuloma and the inflamed alveolar tissue in PTB 22.2. A compressed alveolar zone contains CD163+ macrophages adjacent to granuloma structures (E). Immunofluorescence imaging was performed using CosMx™ spatial profiling with a human protein marker panel including CD163 (red), CD19 (green), CD3 (yellow), CD68 (blue), and vimentin (gray) in panels (C, E). H&E staining was applied in panels (A, B, D, and F). Scale bars: 100 µm (A, B); 200 µm (C–F).

### Supplemental Figure 12

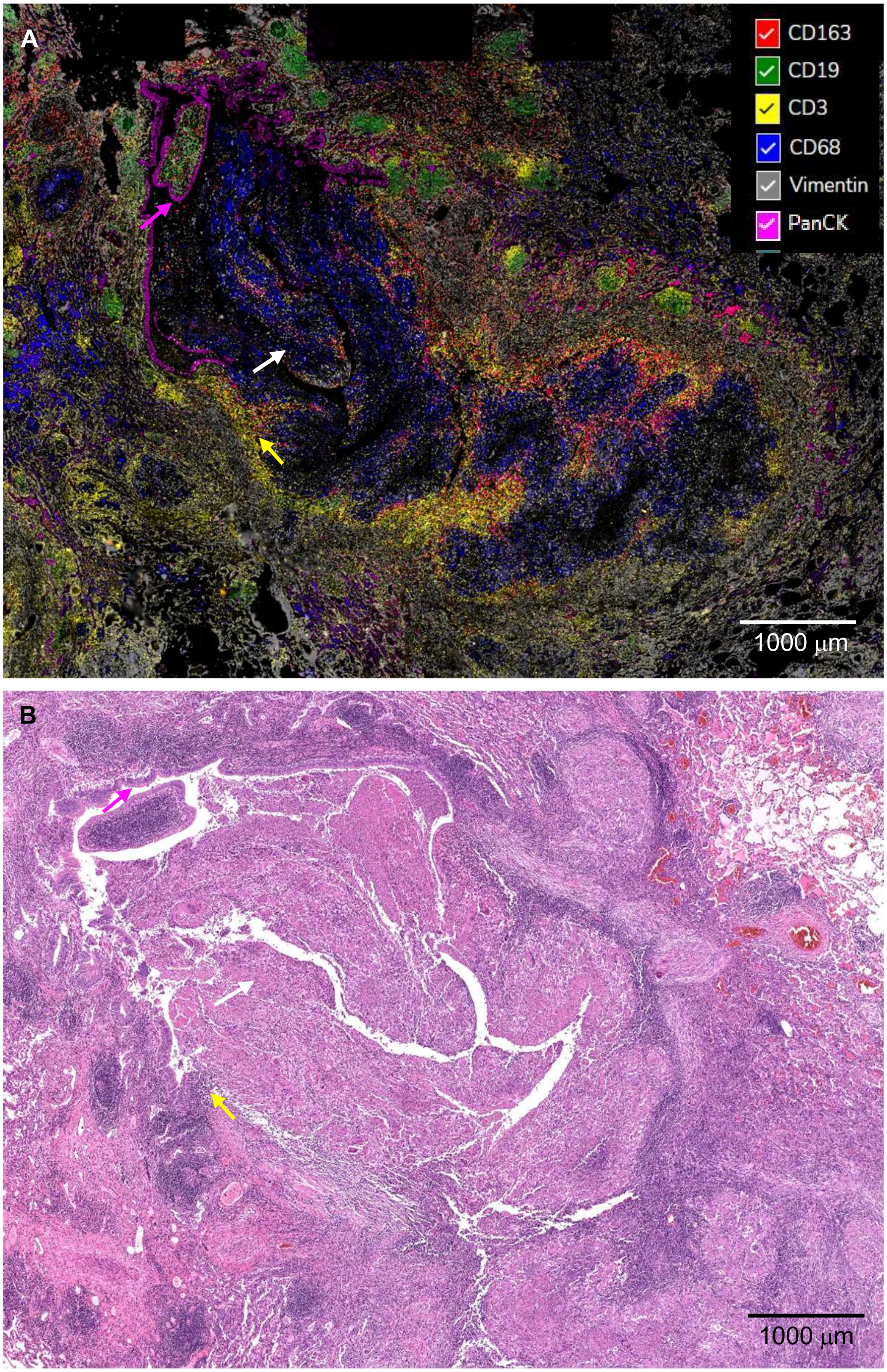


?

**Supplemental Figure 12**. Granulomatous inflammation with necrosis and invasion of the bronchus in pulmonary TB with bronchial obstruction. (A) Immunofluorescence images obtained using the CosMx spatial profiling with the human protein panel, showing markers CD163 (red), CD19 (green), CD3 (yellow), CD68 (blue), vimentin (gray) and PanCK (purple). PanCK is highlighted the bronchial epithelium. (B) Hematoxylin and eosin (H&E) staining of the same region. PTB 22.2 specimen. The epithelium (pink arrows), T cell inflammation (yellow arrows) and areas of necrosis (white arrows) are shown. Scale bars:1000 µm

### Supplemental Figure 13

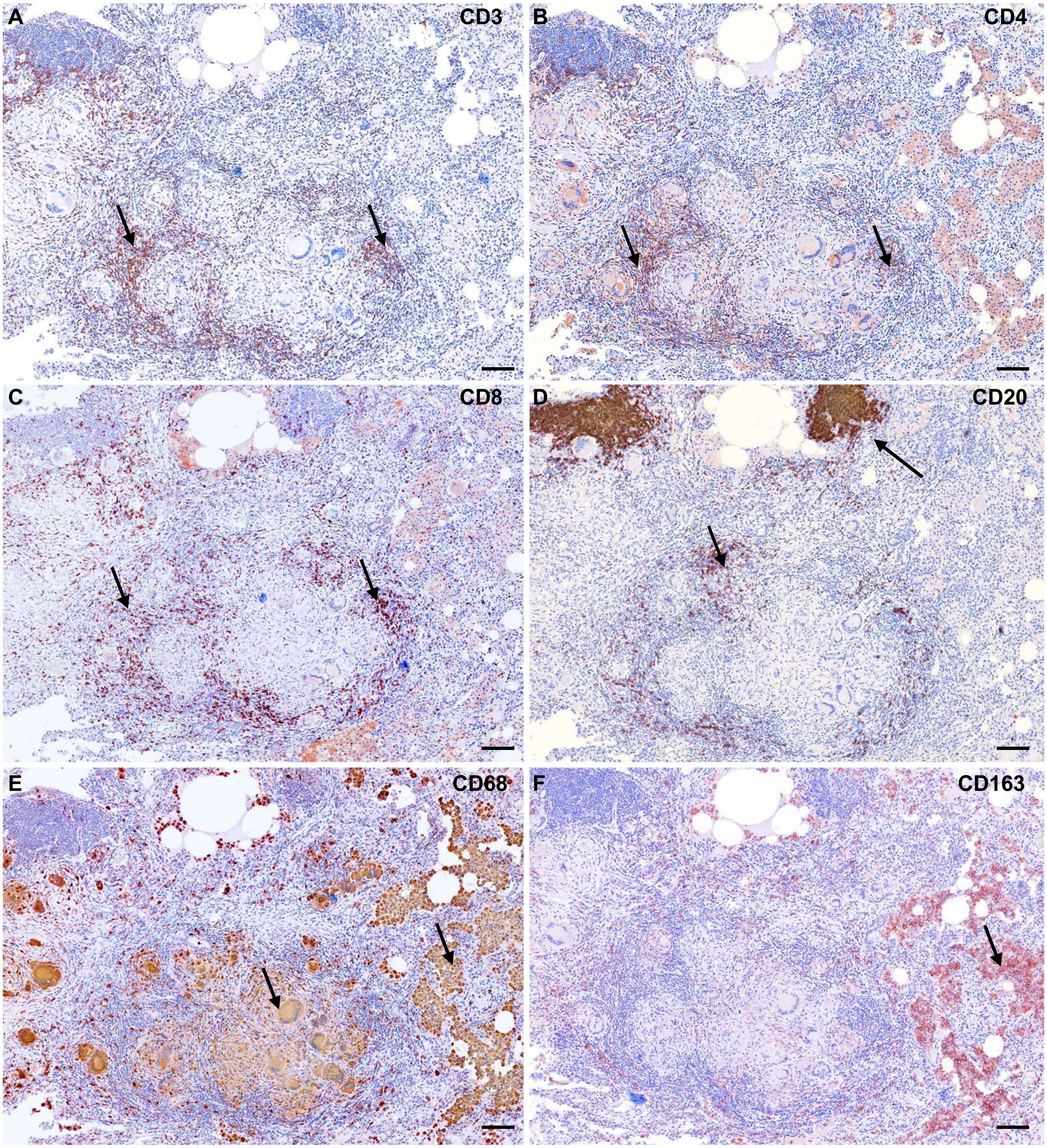


**Supplemental Figure 13**: Immunohistochemical detection of immune cell markers in pulmonary tuberculosis (TB). T cells, including CD3+ (A), CD4+ (B), and CD8+ (C), are predominantly localized to the mantle zone of granulomas. B cells (CD20+) (D) are strongly detected in inflammatory infiltrates outside the granulomas, with some presence in the mantle zone. CD68+ macrophages (E) are abundant in the granuloma core and alveolar regions surrounding the granulomas, while CD163+ macrophages are mainly detected in alveolar areas (F). IHC images are representative of 4 PTB samples. The black arrows show positive cells for each marker, scale bars: 100 µm.

#### Supplemental Figure 14

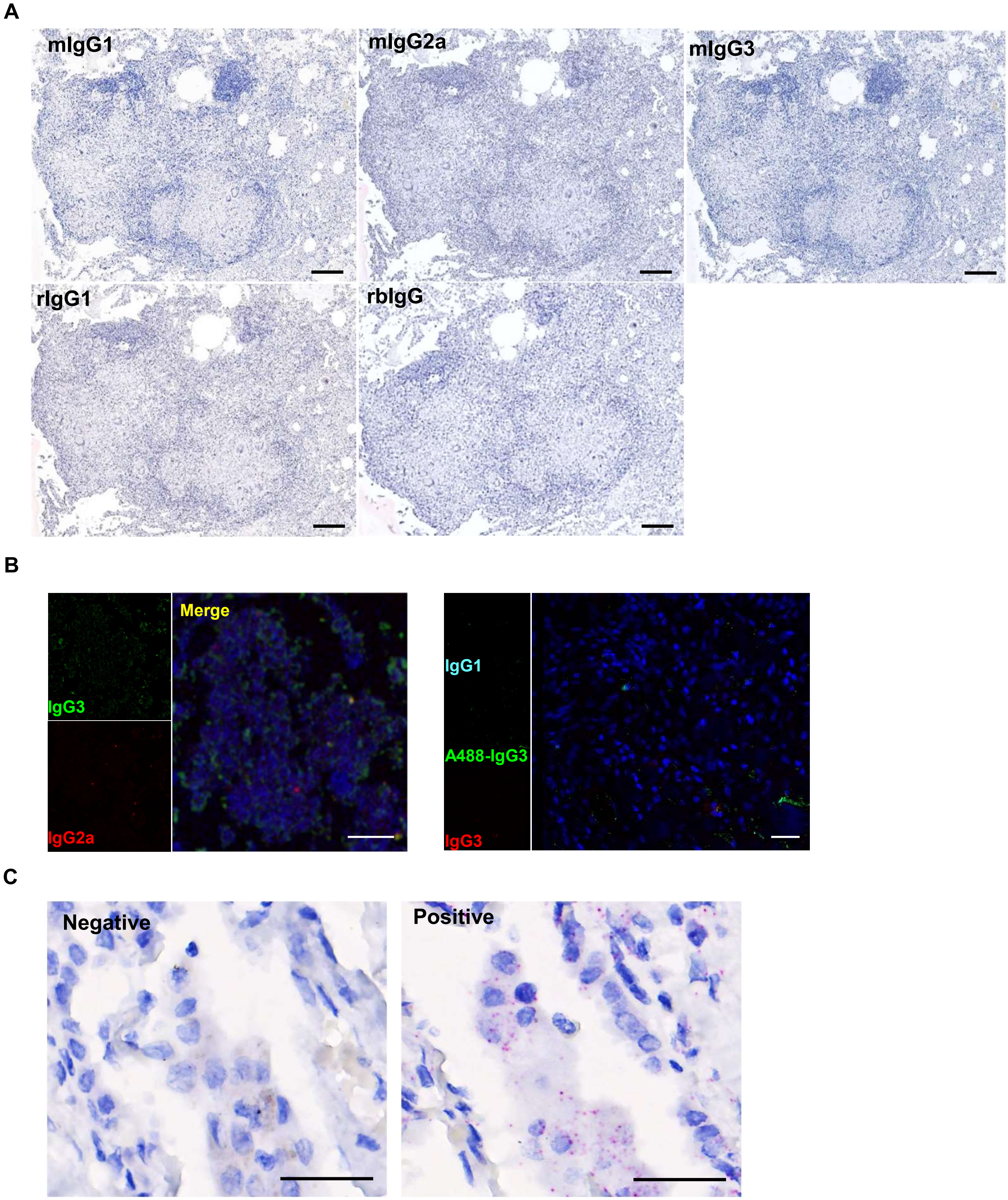


**Supplemental Figure 14.** (A) Negative isotype controls: mouse IgG1 (mIgG1) isotype control for APOE,CD68, CD8, PLIN2 and SPP1; mouse IgG2a (mIgG2a) isotype control for CD20 and CD163; mouse IgG3 (mIgG3) isotype control for TREM2 and TB-LAM; rabbit IgG (rbIgG) isotype control for CD4 and TB-protein and rat IgG1 (rIgG1) isotype control for CD3. (B) Immunofluorescence isotype controls. (C). RNAscope's negative and positive probe controls for foamy macrophages. Negative control images are representative of 4 PTB samples. Scale bars: A, 100; B and C, 20 μm.

#### Supplemental Figure 15

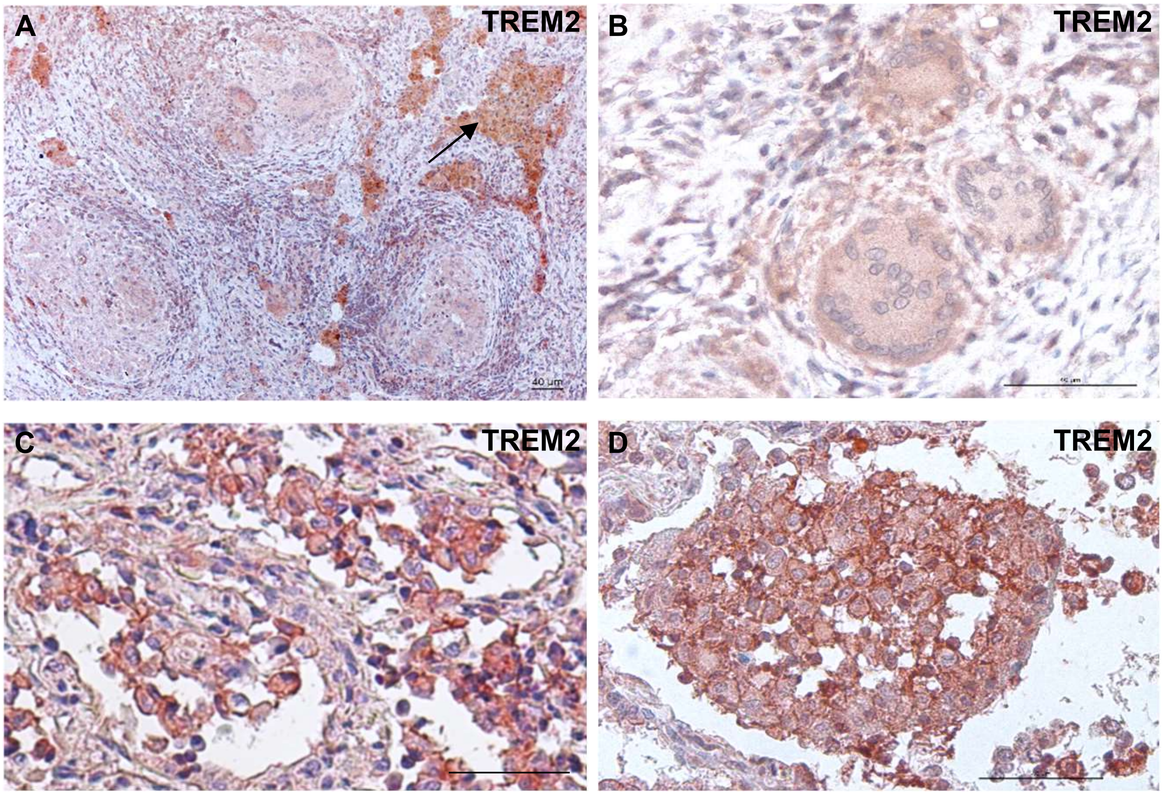


**Supplemental Figure 15**. (A) Detection of TREM2 in the PTB lesion, TREM2 expression is weak in the multinucleated Langhans giant cells (B) and strong in the alveolar foamy macrophages (C-D). IHC images are representative of 4 PTB samples**.** Scale bars: 40 µm (A and D).

#### Supplemental Figure 16

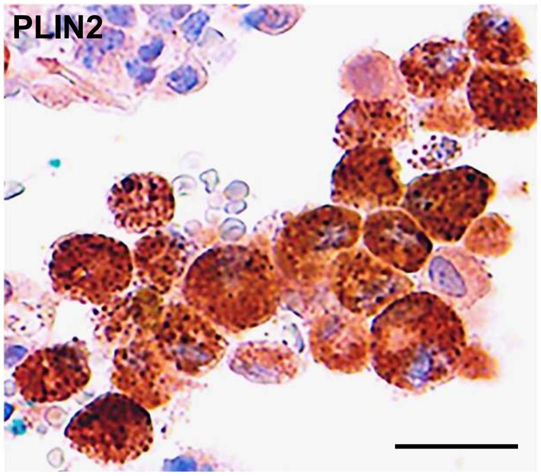


**Supplemental Figure 16. (**A) Alveolar foamy macrophages displaying PLIN2 puncta, indicating the distribution of lipid droplets.

#### Supplemental Figure 17

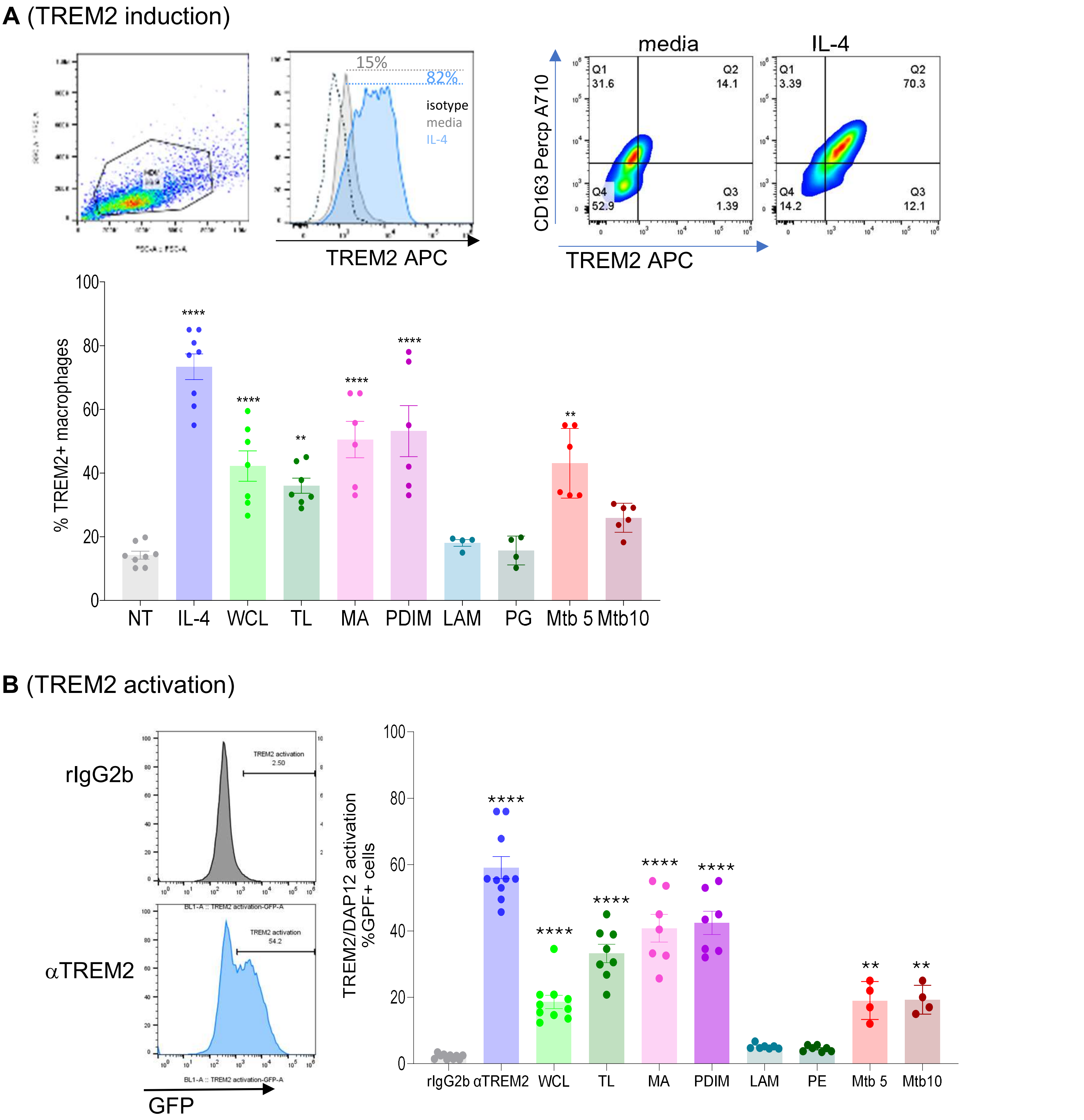


**Supplemental Figure 17.** (A) Human monocyte-derived macrophages (MDMs) were treated with IL-4 (100 ng/mL) and various *M. tuberculosis* (*Mtb*) ligands. CD163 and TREM2 receptor expression were assessed by flow cytometry after 2 days. The graph displays the percentage of TREM2-expressing cells in response to each stimulus (N ≥ 4). (B) GFP-TREM2 reporter 2B4 cells were treated for 24 hours with an anti-TREM2 monoclonal antibody (αTREM2), isotype control (rIgG2b), and various *Mtb* ligands. GFP-positive cells, indicating TREM2/DAP12 activation, were detected by flow cytometry. The graph represents the percentage of GFP+ cells (N ≥ 4). *Mtb* ligands: whole cell lysate (WCL), total lipids (TL), mycolic acids (MA), phthiocerol dimycocerosates (PDIM), lipoarabinomannan (LAM), peptidoglycan (PE), live *Mtb* at MOI 5 (Mtb5), and live *Mtb* at MOI 10 (Mtb10). Dot plots and histograms are representative (n ≥ 5). One-way ANOVA with Tukey multi-comparison test. Statistical significance: ** $p < 0.01$, ***$p < 0.001$, ****$p < 0.0001$.

### Supplemental Figure 18

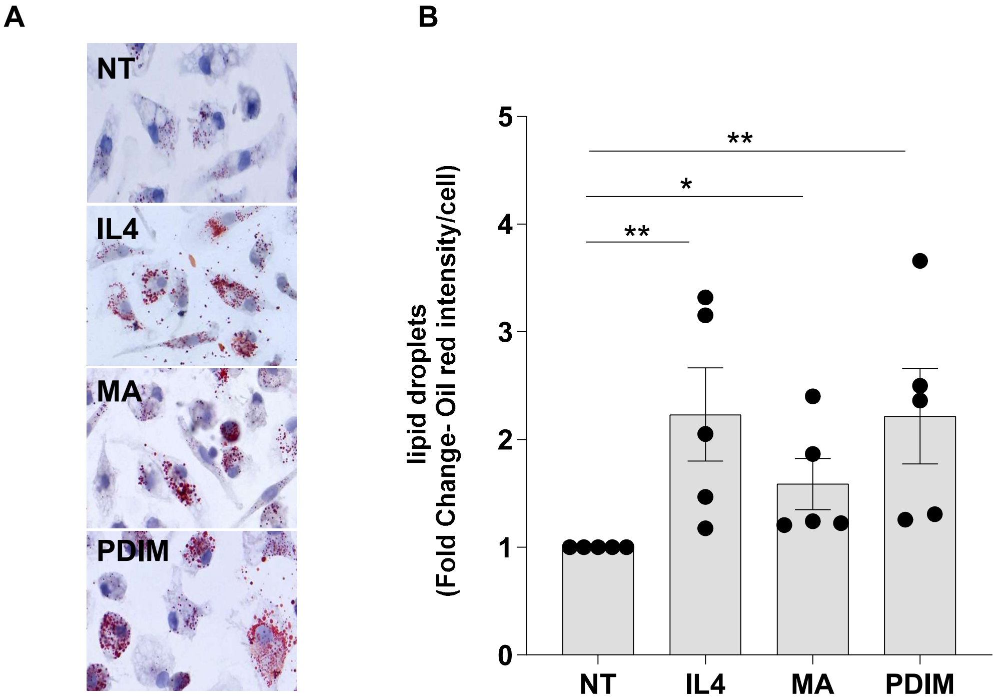


**Supplemental Figure 18:** PDIM and mycolic acids induce lipid droplet accumulation in human monocyte-derived macrophages, similar to IL-4–induced TREM2⁺ macrophages. (A) Representative Oil Red O staining of MDMs under each condition: non-treated (NT), IL-4 (to generate TREM2⁺ macrophages), mycolic acids (MA), and PDIM. (B) Quantification of lipid droplet abundance (fold change in Oil Red –positive droplets per cell) across treatments (n = 5 donors). Both PDIM and MA increase lipid droplet accumulation, with PDIM reaching levels comparable to IL-4–induced TREM2⁺ macrophages. Statistical analysis: one-way ANOVA with uncorrected Fisher's LSD for multiple comparisons. Statistics were performed on raw lipid droplet counts prior to normalization; fold-change values are displayed for visualization. Significance: $p < 0.05$; $p < 0.01$.

### Supplemental Figure 19

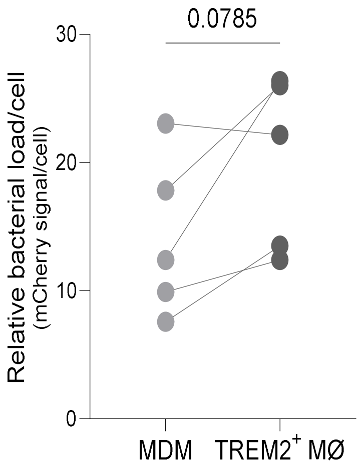


**Supplemental Figure 19.** Macrophages were infected with *Mtb* GFP-mCherry, with inducible GFP and constitutive mCherry expression (n = 5). Images were acquired using confocal microscopy, with DAPI used to stain nuclei. Quantification of constitutive mCherry was performed using ImageJ. One-way ANOVA with Tukey multi-comparison test. Statistical significance: * $p < 0.05$.

#### Supplemental figure 20

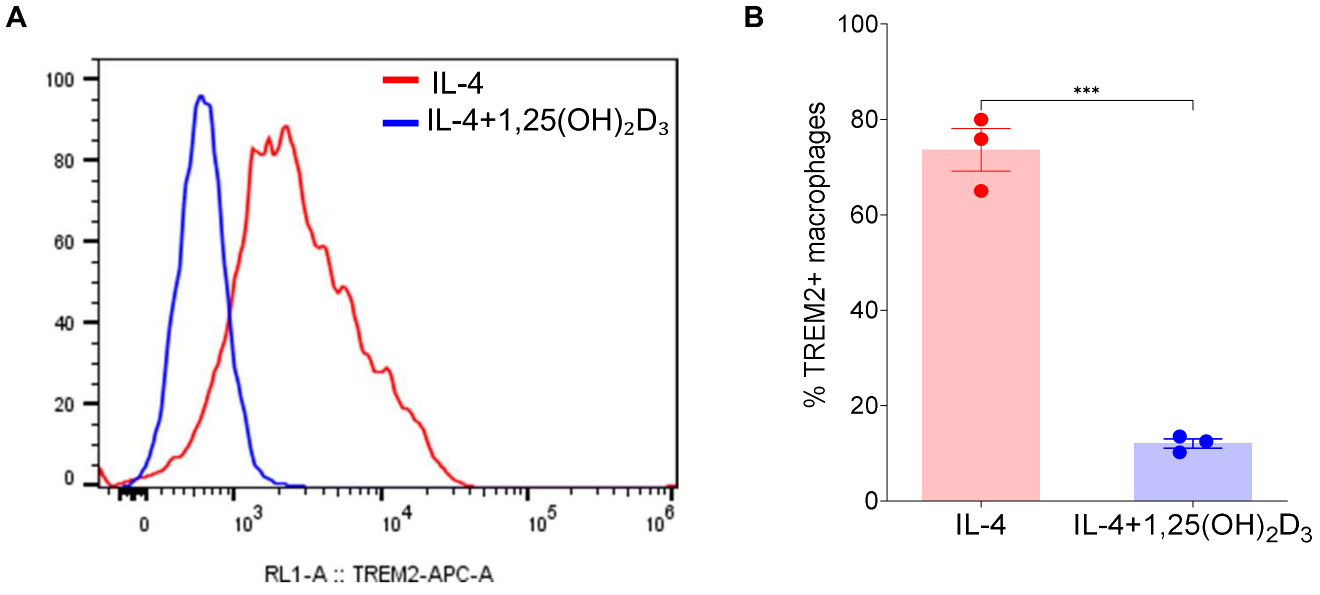


**Supplemental Figure 20. (**A) TREM2⁺ macrophages were treated with 1,25-dihydroxy-vitamin D3 ($1,25(OH)_2D_3$). TREM2 receptor expression was assessed by flow cytometry after 1 day. The graph displays the percentage of TREM2-expressing cells in response to each stimulus (N = 3). The histogram is representative of a total of 3 donors. (B) One-way ANOVA with Tukey multi-comparison test. Statistical significance: *** $p \leq 0.001$.

### Supplemental Figure 21

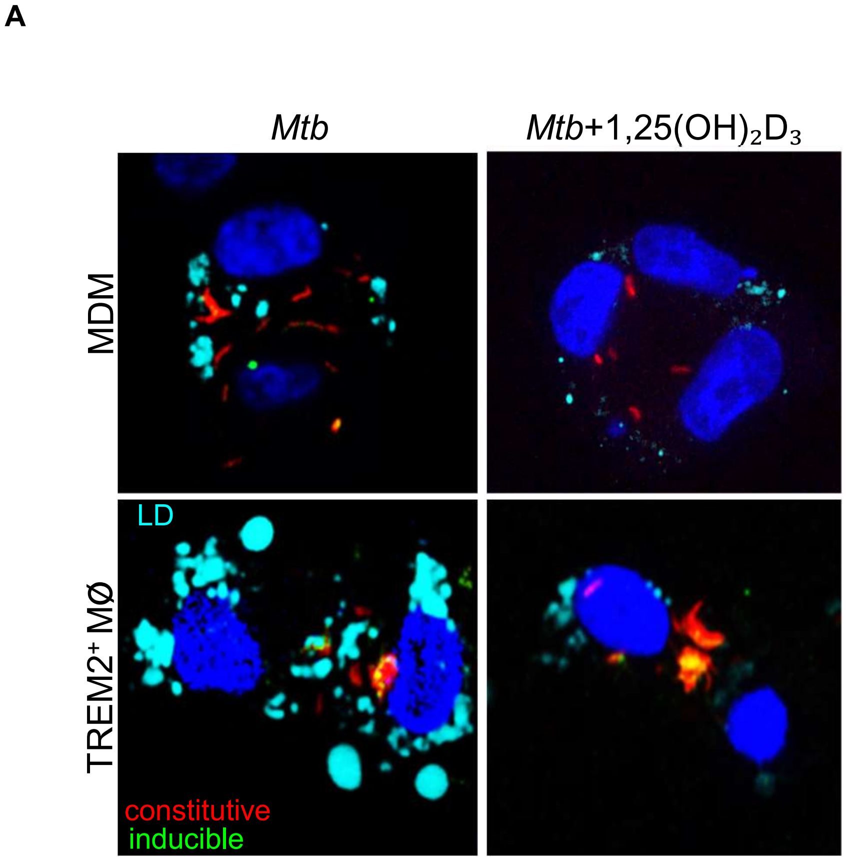


**Supplemental Figure 21.** Lipid droplets and *Mtb* viability are quantitatively linked in MDMs and TREM2⁺ MØ. (A) Representative confocal images of MDMs and TREM2⁺ macrophages infected with *Mtb* ± 1,25$(OH)_2D_3$, showing lipid droplets (SMCy5.5-cyan) and Mtb reporter signal (mCherry-constitutive/GFP-inducible). TREM2⁺ MØ display elevated LD accumulation that is reduced by 1,25$(OH)_2D_3$. N=4

**Supplemental Figure 22**

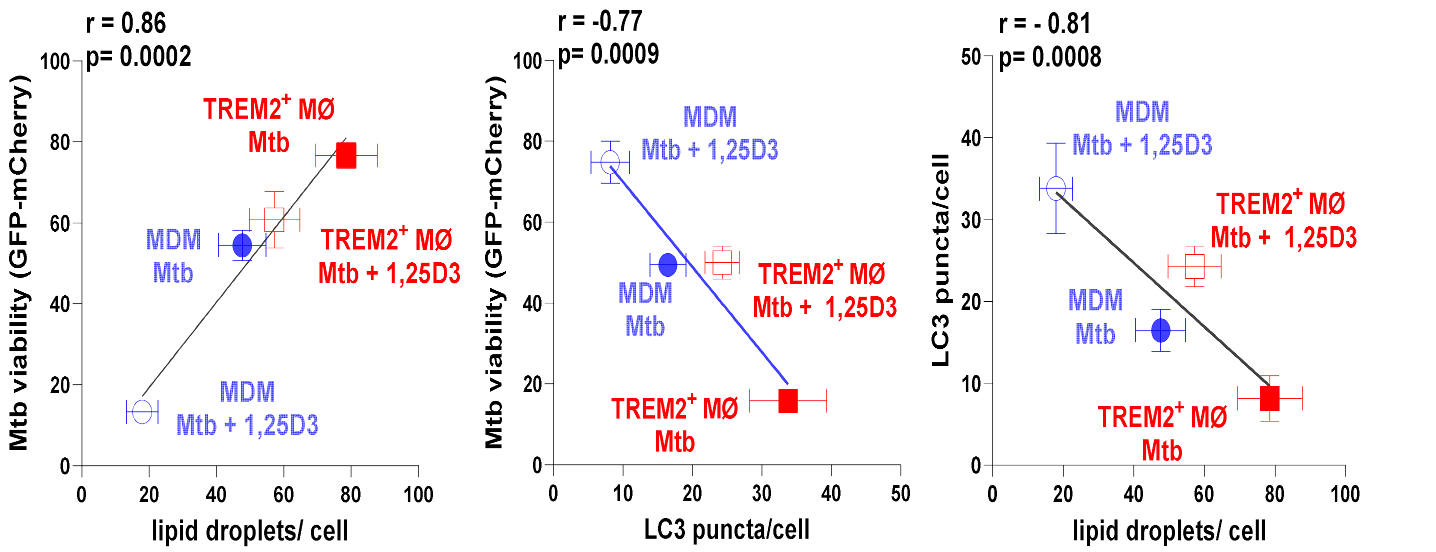


**Supplemental Figure 22**. Pearson correlation analysis of lipid accumulation, autophagy, and bacterial viability. Donor-matched analyses performed side by side using the same experimental samples comparing unpolarized monocyte-derived macrophages (MDM, blue) and TREM2$^+$ macrophages (red) following Mycobacterium tuberculosis infection, with or without 1,25-dihydroxyvitamin $D_3$ (1,25D3). Each symbol represents one donor. Pearson correlation analyses show positive associations between lipid droplet abundance and bacterial viability and inverse associations between LC3 puncta formation and bacterial metabolic activity. Correlation coefficients (r) and two-tailed p values were calculated as described in Methods.
